# Intracortical brain-computer interface for navigation in virtual reality in macaque monkeys

**DOI:** 10.1101/2025.05.05.652199

**Authors:** Ophelie Saussus, Sofie De Schrijver, Jesus Garcia Ramirez, Thomas Decramer, Peter Janssen

## Abstract

We present an innovative intracortical Brain-Computer Interface (BCI) to bridge the gap between laboratory settings and real-world applications. This BCI approach introduces three key advancements. First, we utilized neural signals from three macaque brain regions – primary motor, dorsal and ventral premotor cortex – enabling precise and flexible decoding of real-time three-dimensional (3D) sphere/avatar velocities. Second, we developed a realistic, immersive 3D virtual reality setup with dynamic camera tracking, allowing continuous navigation and obstacle avoidance that closely mimic real-world scenarios. Finally, our BCI approach is very well suited for use by paralyzed patients, featuring a brief passive fixation without overt movements and closed-loop operation without retraining of the decoder during online decoding, relying on the user’s neural plasticity and the decoder’s robust generalization across tasks. Our BCI adapted to different environments, targets, and obstacles, illustrating its potential to substantially enhance the quality of life for paralyzed patients by enabling natural, reliable and flexible control in complex settings.

## Introduction

Brain-Computer Interfaces (BCI) are a promising technology that enable direct communication between the brain and external devices by translating neural activity into specific commands. BCIs can be used to enhance or restore functions lost due to disease or trauma, bypassing damaged biological pathways(*1–3*). Intracortical BCIs (iBCIs) in nonhuman primates and humans have enabled control of computer cursors(*4–11*), manipulation of robotic or prosthetic arms(*6*, *10–17*), wheelchair control(*18*, *19*), restoration of communication(*20–24*), restoring motion of paralyzed limbs(*25–27*), and even the provision of sensory feedback(*28*).

Previous motor BCI studies have often focused on cursor or robotic arm control due to the limitations of a lab environment, which hamper the translation to real-world situations such as wheelchair navigation. Furthermore, realistic brain-controlled real-world navigation is frequently confronted with unpredictable events requiring highly flexible online corrections that are likely to depend on both premotor and primary motor cortex activity. To bridge the gap between lab and home environments, we developed a BCI approach in a 3D virtual reality (VR) environment with stereoscopic vision for realistic simulations of increasing complexity and unpredictability. To maximize the potential of flexible control of the cortical motor system, we employed for the first time simultaneous recordings in dorsal and ventral premotor and primary motor cortex. While previous BCI research was limited to simple tasks in basic settings (e.g., a sphere moving freely in a featureless space)(*8*, *29*), our approach offers a more complex and realistic VR environment, with 3D movements that closely mimic real-life scenarios. Additionally, we implemented a navigation task that effectively mimicked real-life wheelchair control through dynamic camera tracking and a realistic 3D VR environment.

Our iBCI successfully translated neural activity from three motor cortical areas into real time velocities in three monkeys, demonstrating the potential of our BCI system for real-world brain-controlled navigation.

## Results

We implanted three rhesus monkeys with Utah arrays in three motor regions: primary motor cortex (M1), dorsal premotor cortex (PMd) and ventral premotor cortex (PMv). We designed five tasks to validate the effectiveness of our BCI (Table 1). In all tasks, the decoder was trained with data obtained during passive observation of the movements in the VR environment without any overt movements.

**Table 1.** Estimated chance level, success rate (mean ± standard deviation), number of channels used for decoding (mean ± standard deviation) and the number of sessions per task and per monkey. For Monkey 1, the average percentages of electrodes used for decoding in M1, PMv and PMd were 46%, 34% and 20% respectively. For Monkey 2, only electrodes in M1 were used. For Monkey 3, the percentages of electrodes used for decoding in M1, PMv and PMd were 36%, 41% and 23% respectively. The smaller percentage in PMd in monkeys 1 and 3 was because only 64 electrodes were available for recording, compared to 96 for M1 and PMv.

| Task | Monkey | Chance level [%] | Success rate* [%]<br>(mean ± std) | Number of channels<br>(mean ± std) | Number of sessions |
| --- | --- | --- | --- | --- | --- |
| Center-out task | Monkey 1 | 23 | 73 ± 5 | 140 ± 6 | 5 |
| | Monkey 2 | 20 | 83 ± 8 | 49 ± 3 | 10 |
| | Monkey 3 | 23 | 64 ± 13 | 217 ± 11 | 19 |
| Continuous Navigation task | Monkey 1 | 34 | 69 ± 6 | 139 ± 6 | 6 |
| | Monkey 2 | 30 | 70 ± 15 | 47 ± 5 | 7 |
| | Monkey 3 | 33 | 59 ± 8 | 216 ± 10 | 15 |
| 3D Center-out task | Monkey 2 | 17 | 69 ± 6 | 43 ± 5 | 12 |
| | Monkey 3 | 19 | 57 ± 8 | 202 ± 14 | 10 |
| Respawn task | Monkey 2 | 24 | 76 ± 9 | 40 ± 4 | 11 |
| | Monkey 3 | 26 | 82 ± 5 | 199 ± 15 | 9 |
| Obstacle task | Monkey 2 | 19 | 63 ± 6 | 40 ± 4 | 10 |
| | Monkey 3 | 23 | 69 ± 9 | 203 ± 17 | 11 |

\*All success rates significantly exceeded chance level ( $p < 0.0001$ , permutation test)

### Success rates

To assess how well macaque monkeys could perform BCI-control tasks in a moderately complex 3D environment in VR, we first implemented the Center-out and Continuous Navigation tasks (Supplementary Movies S1 and S2). All three monkeys performed significantly above chance level from the start of the recording period (Table 1, permutation tests: all p < 0.0001), with individual sessions achieving success rates as high as 96% (Fig. 1A, Fig. 1B and Fig. S1). Performance in the Continuous Navigation task was slightly lower (4 – 13% fewer successful trials) compared to the Center-out task, most likely due to the dynamic camera tracking in the former task. Monkeys 1 and 2 had participated in a previous BCI project and reached success rates around 70% from the start of the recording period in both tasks. In contrast, Monkey 3 required more sessions (N=11 for the Center-out task and N = 14 for the Continuous Navigation task) to achieve a similar success rate, demonstrating a clear learning trend across sessions in the Center-out task (the first task learned; linear regression: β=0.0156, F(1,17)=12.98, p=0.0022, coefficients of determination (*R^2^*)=0.433). To verify that navigation was also possible from a first-person perspective, we removed the monkey avatar in the Continuous Navigation task in 3 sessions. Monkey 3 achieved similar success rates (average success rate of 54±1%) as in the standard Continuous Navigation task, confirming the adaptability of the BCI across perspectives (Supplementary Movie S3). Thus, macaque monkeys are able to achieve accurate navigation in a 3D VR environment, even with dynamic changes in camera viewpoint during navigation, as well as from a first-person perspective.

**Fig. 1.**
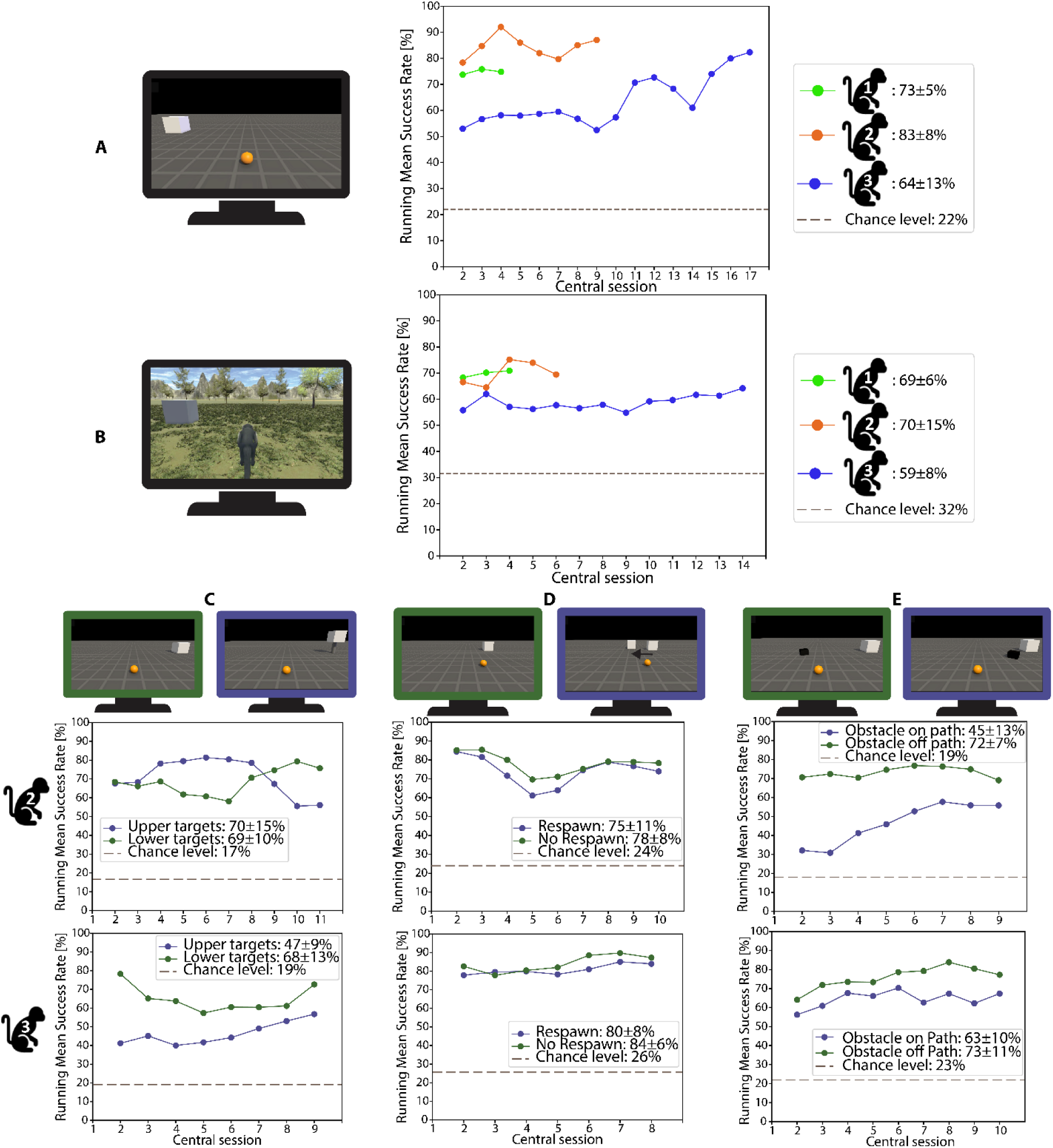
BCI control performance and flexibility across tasks and conditions. **A-B** Running mean of the success rate across sessions (window size = 3 and sliding window = 1), shown together alongside the estimated chance level (dashed line). The average success rate ± standard deviation across all sessions is also displayed. **A** Running mean success rate of Monkey 1, 2 and 3 for the Center-out task. **B** Same data for the Continuous Navigation task. **C-D** Performance breakdowns demonstrating flexibility of BCI control in more complex conditions. Each panel shows running mean success rate for Monkey 2 (top) and Monkey 3 (bottom) under two conditions per task: **C** 3D Center-out task with upper (blue) and lower (green) targets. The success rates for reaching the upper and lower were nearly identical (69 and 70, respectively) and well above chance level for Monkey 2. Performance in this task was lower for Monkey 3, but the monkey still performed well above chance level for both targets. **D** Respawn task with trials that included respawning (blue) and no respawning (green). **E** Obstacle task with the obstacle on path (blue) and off path (green).

To further investigate to what extent our animals could achieve flexible and accurate BCI control in more challenging VR environments, we later designed three additional tasks, which were performed in separate sessions. In the 3D Center-out task (Fig. 1C, Supplementary Movie S4), in which the animal was required to move the sphere in three dimensions, the overall success rate averaged 69±6% across all sessions for Monkey 2 and 58±8% for Monkey 3. Although performance was lower than in the 2D Center-out task, it remained significantly above chance level from the start of the recording period (Table 1, permutation test: p < 0.0001), which is expected given the increased difficulty of controlling the movement in three dimensions. Both monkeys successfully controlled movements in the third dimension (z-direction) without requiring additional behavioral training beyond the standard Passive Fixation phase for this task, suggesting a degree of intuitive adaptation to 3D control.

We also observed markedly flexible and accurate BCI control in the Respawn task (Fig. 1D, Supplementary Movie S5). Without the need for explicit decoder training or additional animal training, both monkeys tested achieved around 90% success when the target object unpredictably shifted position during the trial with no significant difference in performance between respawn trials and no-respawn trials (Mann-Whitney U = 185629.5, p = 0.1556 for Monkey 2 and U = 101694.5, p = 0.0765 for Monkey 3).

Both monkeys also learned to achieve reliable obstacle avoidance in the Obstacle task (Supplementary Movie S6). Initially, the success rate of Monkey 2 was lower in trials with an obstacle on the path (36%) compared to off path trials (66%, Fig. 1E), but at the end of the recording period, the success rate of this animal in on path trials reached 55% (which was still significantly lower than in off-path trials, U = 142032.5, p = 7.071 × 10^-16^). In contrast, Monkey 3 performed markedly better in on-path trials, achieving an average success rate of 63%, closer to its off-path performance (73%, U = 132242.5, p = 6.534 × 10^-4^, Fig. S1E). Overall, the monkeys successfully completed the various tasks with a high success rate and without the need for extensive training, even when navigating 3D trajectories, encountering obstacles, or adjusting to changes in target position during the trials.

### Decoder generalization

Our decoder exhibited robust generalization across tasks, target positions, and environments. It was trained during a brief Passive F phase (∼7 minutes), in which the monkey passively observed movements toward three targets in a standard Center-out layout (Fig. S2A) with no obstacles or dynamic events. This single decoder was then used — without retraining or recalibration — to decode neural activity online in three distinct tasks: the Center-out, Obstacle (featuring object avoidance), and Respawn tasks (involving mid-trial target jumps). This fixed-decoder approach capitalizes on neural adaptation and allows to assess decoder robustness across varying task demands.

The decoder also generalized to novel target positions not included during training. To test this, we compared success rates between pretrained targets shown during training (left, straight, right) and novel ones (slight left, slight right). Across all tasks and monkeys, performance on novel targets was comparable to — and in some cases significantly exceeded — that on pretrained targets (Fig. S3). The higher performance on new targets likely reflects differences in task difficulty: pretrained targets required larger movement amplitudes, while novel targets were closer to the origin. Importantly, success rates for all targets remained well above chance, demonstrating effective generalization of the decoder and behavioral strategy.

The decoder also generalized across environments and agent representations. In the Continuous Navigation task, although the decoder was trained on passive observations of a sphere moving on a simple grid with dynamic camera tracking (Supplementary Movie S7), it supported accurate online control in a visually distinct forest environment without any training in that setting. Furthermore, decoder performance remained stable across different visual representations of the controlled entity — including a sphere, a monkey avatar, and a first-person perspective. These results suggest that the decoder captured behaviorally relevant neural dynamics while remaining robust to changes in visual and contextual features, supporting its potential usability in real-world settings.

In addition to across-task generalization, we assessed whether monkeys could adapt within a single session and in a given task under fixed-decoder conditions. To assess whether monkeys improved control with a fixed decoder within a single session, we applied the within-session performance trend analysis described in the Methods (Fig. S4). For the initial tasks (Center-out and Continuous Navigation), 37% of sessions showed significant within-session improvement, 20% a decline, and 43% showed no change (constant). In contrast, in more difficult tasks, performance improvement occurred in only 20% of sessions, while 27% showed a decline (for data on individual monkeys, see Figure S4B). These findings highlight variability in short-term performance changes across tasks and subjects, which may reflect differences in the ability or tendency to adapt neural activity during BCI control.

### Offline decoding reveals regional contributions and neural adaptation

To better understand the contributions of each brain area to decoder performance, we performed offline decoding using a matched number of high-weight channels from each area in the two animals (Monkey 1 and 3) that had spiking activity in the three areas. Decoding success rate was assessed using neural signals from individual areas (M1, PMd, PMv), pairwise combinations (PMv+PMd, M1+PMd, M1+PMv), an equal-area “All” condition (PMv+PMd+M1), and an “Online simulation” condition using all available channels offline. These success rates were compared to the “Online decoding” performance achieved during real-time, closed-loop control with visual feedback. Across both monkeys and tasks, PMv and PMd consistently outperformed M1 (Fig. 2A–B), with several comparisons reaching statistical significance (see Table S1). The PMv+PMd combination yielded the highest performance among area pairs, significantly outperforming M1+PMd and M1+PMv (all p<0.05 in both monkeys, except Continuous Navigation task Monkey 3, see Table S1). These results indicate that PMv and PMd signals are synergistic, and that M1 contributes little additional independent information once premotor activity is included. In the Center-out task, PMv+PMd achieved performance statistically indistinguishable (p>0.05) from both the full All condition and the Online simulation, demonstrating that premotor signals alone can support high BCI performance. In the Continuous Navigation task, slight performance advantages of the full-channel configurations may reflect differences in total channel count rather than meaningful additional contributions from M1.

**Fig. 2.**
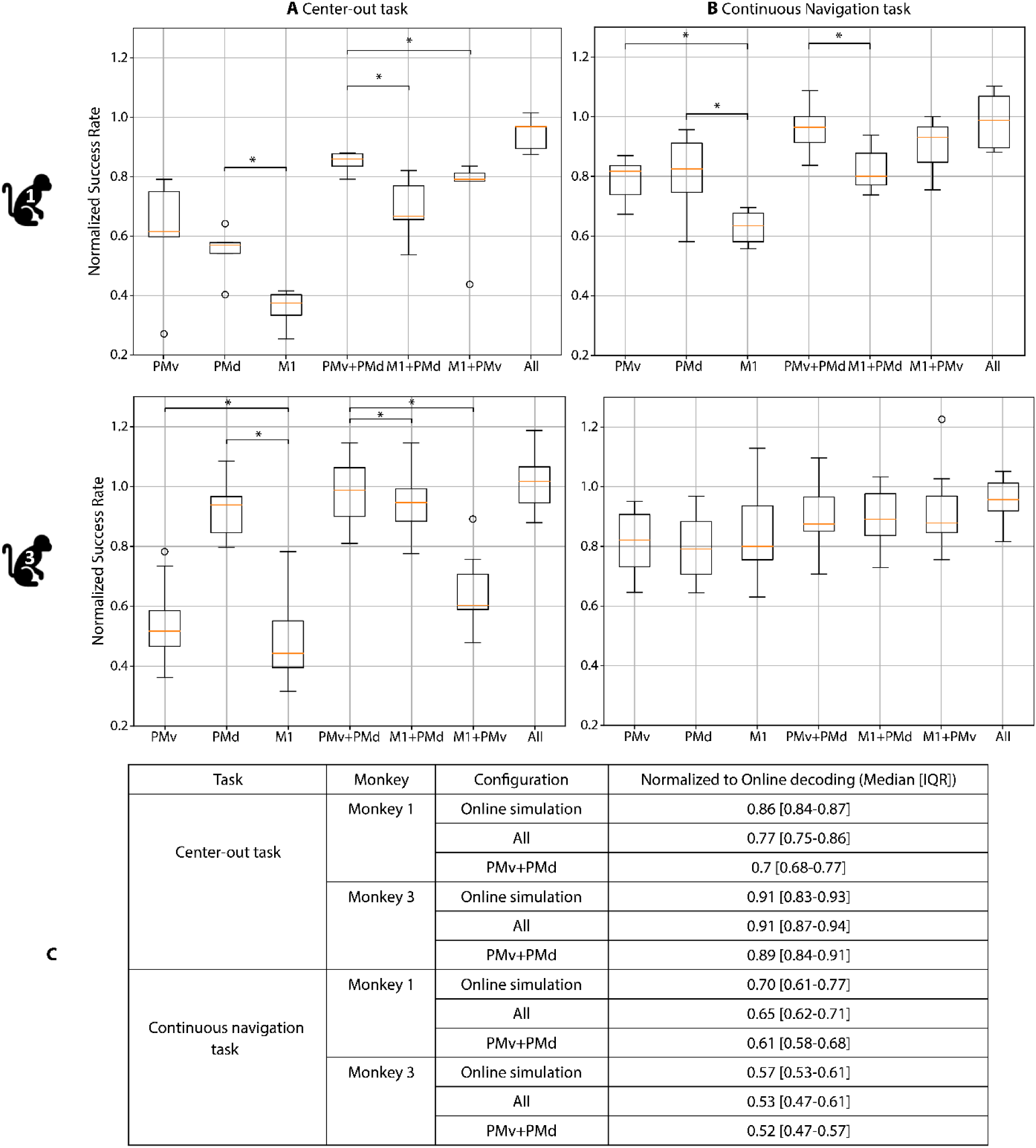
Offline decoding performance across cortical regions and configurations, compared to both offline simulation and actual closed-loop online performance. **A, B** Offline decoding success rates for different cortical input configurations in the **A** Center-out task and **B** Continuous Navigation task, using data from Monkeys 1 and 3. Each configuration consisted of a matched number of high-weight electrodes from either individual areas (PMv, PMd, M1), pairwise combinations (PMv+PMd, M1+PMv, M1+PMd), or an equal-area “All” condition (balanced channels from each area). Success rates were normalized to the “Online simulation” condition (decoder trained and tested offline using all available channels from that session). Boxplots show the median (orange line), interquartile range (box), and range within 1.5× IQR (whiskers); circles indicate outliers. Asterisks mark significant differences within each monkey (Monkey 1: one-sided exact binomial test; Monkey 3: Wilcoxon signed-rank test; p < 0.05). All statistical comparisons and p-values are provided in Table S1. **C** Table summarizing median success rates [IQR], normalized to the true Online decoding performance (closed-loop condition with real-time feedback and adaptation). IQR computed from empirical session medians. Offline decoding underperforms across all configurations, highlighting the added value of neural adaptation during real-time BCI control. Number of sessions (N) for each task is provided in Table 1.

These decoding patterns closely aligned with the decoder weight analysis, where the majority of weight was assigned to PMv and PMd electrodes. In Monkey 1, decoder weights were distributed as 60% PMv, 21% PMd, and only 19% M1. In Monkey 3, the pattern shifted slightly (42% PMd, 19% PMv, and 39% M1), but premotor areas still dominated. In monkey 2—who performed worse on the Obstacle Task—all electrodes used in the decoder were from M1 since the other arrays had no electrodes with spiking activity. These observations suggest that premotor areas may be important for successful obstacle avoidance.

Importantly, decoding performance in all offline conditions remained consistently lower than the actual online performance (Fig.2C), even when using the same decoder and neural data (Online simulation). This performance gap points to the presence of neural adaptation during closed-loop control. Consistent with this interpretation, our within-session trend analysis showed that Monkey 1, who exhibited the largest gap between online and offline decoding, also had the highest proportion of sessions with performance improvement over time (53.4%), indicating dynamic adjustment of neural activity during task execution (Fig. S4B). These findings provide strong evidence that monkeys adapt their neural output in response to decoder feedback, enhancing control over time. The inability to replicate this effect offline underscores the importance of real-time adaptation mechanisms in iBCI performance.

### Trajectories and time to target

We investigated the sphere’s trajectories during BCI control in the additional tasks. Fig. 3A shows that in both monkeys, the final y-coordinate of the sphere in the 3D Center-out task differed significantly between upper and lower target trials (Mann-Whitney U = 195291, p = 5.26×10^-60^ for Monkey 2; U = 75468, p = 3.11×10^-34^ for Monkey 3).

**Fig. 3.**
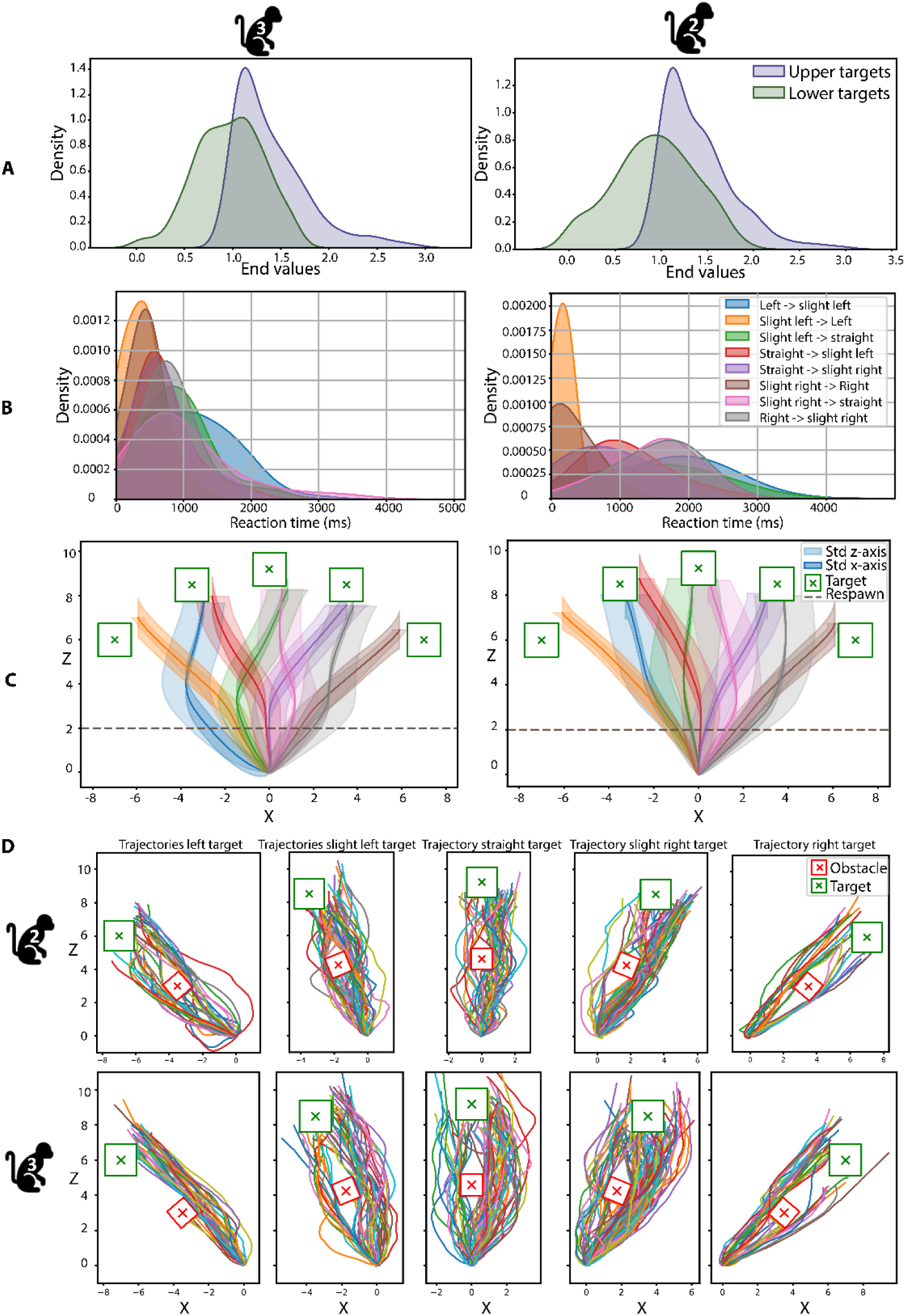
Trajectories analysis of the three supplementary tasks for Monkeys 2 and 3. **A** 3D Center-out task: Density distribution of the y-coordinate of the endpoint values of trials with upper (blue) and lower (green) targets (across all sessions). **B-C** and Reaction time and average trajectory per respawn condition for the Respawn task. B Density distribution of average reaction time per respawn condition over all trials and sessions. **C** Average trajectory for each respawn condition, together with the standard deviation (Std) in the x- and z-directions across all trials and all sessions. The respawn moment is also shown. **D** Individual trajectories of successful trial across all sessions for the Obstacle task, for each target (green square) when the obstacle (red square) was on the target’s path. The number of sessions (N) for each task is provided in Table 1.

Next, we investigated how fast Monkeys 2 and 3 corrected the trajectories during respawning by calculating the reaction time, defined as the time required for the sphere to change its trajectory from the original target to the respawn target, starting from the moment the target was respawned (i.e. changed location). The average reaction time across all conditions was 661ms for Monkey 2 and 1058ms for Monkey 3 (Fig. 3B and C, Kruskal-Wallis test between conditions, Monkey 2: statistic = 91.84, p = 5.18 × 10^-17^; Monkey 3: statistic = 41.77, p = 5.759 × 10^-7^).

Finally, we explored the monkey’s strategies for obstacle avoidance by plotting the trajectory of each individual trial per target for each monkey (Fig. 3D). Monkey 2 utilized both sides to navigate around the obstacles and made deviations from the straight trajectory closer to the target, whereas Monkey 3 tended to favor one side and often made a larger deviation early in the trajectory.

In the Center-out task, the average time to target in the Passive fixation phase, during which the sphere was autonomously controlled by the Unity 3D engine’s built-in navigation system (see Methods section: “Experimental phases” and “Unity”), was 2750ms. This compares to 3196ms (Monkey 1), 3182ms (Monkey 2), and 3111ms (Monkey 3) in the Online Decoding phase, which was controlled in real time by the monkeys through neural activity. This result indicates that the time to target in the Online Decoding phase was on average 15% slower (across all monkeys) compared to the Passive Fixation phase (Fig. S5A). In the Continuous Navigation task, the average time to target was 48% slower during the Online Decoding phase (Fig. S5B), which highlights the increased difficulty of this task. For the time to target in the three additional tasks, see Fig. S5 C-E (see also Supplementary Results for additional task-specific comparisons).

### Influence of surface electromyography (sEMG) and eye movements on decoded velocities

We wanted to assess the influence of sEMG on the decoded velocities (v, v_x_, v_y_, v_z_). The mean correlation coefficients across sessions between sEMG and decoded velocities were consistently very low across tasks and velocity components, ranging from −0.08 to 0.06 for Monkey 1 and −0.20 to 0.13 for Monkey 2 (Table S2). Cross-correlation analysis for lags ranging from −100 to 100ms (50ms increments) yielded similar results. Regression analyses further supported these findings, with *R^2^* values near zero for both the linear and non-linear models (Table S3). Although the combined p-values from the regression analyses were statistically significant, the small effect sizes suggest that sEMG had minimal influence on decoded velocities. Furthermore, the high mean squared error (MSE) values for both monkeys across all tasks and velocity components indicate poor predictive performance of the models (Table S3).

We also computed the Spearman correlation between the *x* and *y* components of the eye position and the decoded velocity. Similar to the results for the sEMG analysis, we found very small correlation coefficients for all velocity components and for each component of the eye movement, except for a notable correlation between the *x* component of the eye position and *v_x_* (Table S4), which is related to the monkeys tracking the sphere’s position during online decoding, as in natural behavior(*30*). However, such a correlation was not observed during the Continuous navigation task, where the camera dynamically followed the avatar. In this task, the decoding was world-centered rather than eye-centered (decoding velocities to the left remained left even if the camera had moved and the avatar had to go straight ahead), altering the relationship between the eye position and the decoded velocity. These findings suggest that eye movements did not fundamentally influence the decoded velocity.

## Discussion

We developed a robust and flexible iBCI that enables macaque monkeys to control movement in a 3D virtual environment using only neural activity. By decoding signals from M1, PMd, and PMv into continuous velocity commands, our system supported real-time control across a broad range of tasks including center-out navigation, obstacle avoidance, dynamic target switching, and first and third person continuous navigation in visually complex scenes, mimicking real-life navigation. Unlike prior studies that focused on 2D cursor control or robotic arms, our virtual-reality design emphasizes real-world challenges: task switching, spatial and contextual generalization, and BCI control without overt movements or proprioception.

Our decoder introduces key innovations relative to previous iBCIs. To allow a detailed comparison, we have summarized the methodology employed in previous studies and in our study in Table S5. A central innovation of our approach lies in the training paradigm. Our decoder was trained during a single, brief passive observation phase (∼7 minutes) with no overt movements, and was then applied without recalibration or retraining across multiple tasks. This fixed-decoder design stands in contrast to most previous iBCI systems, which relied on overt or attempted movements during training and often included multi-phase training or online adaptation (e.g., ReFIT(*31*)). In our case, decoder training was based entirely on passive visual feedback, and control was achieved using neural signals alone, under full physical restraint, with sEMG verification to rule out residual movement correlation. These constraints closely replicate the conditions faced by individuals with complete paralysis.

Our decoder also differs architecturally from standard approaches. Instead of directly mapping neural activity to movement using linear models (e.g., Kalman or Wiener filters), we adapted the PSID framework for online, closed-loop decoding, and replacing the linear regression with a nonlinear variant. Our method extracts a low-dimensional latent state optimized for behavioral relevance before predicting velocity, offering a favorable trade-off between complexity, interpretability, and real-time feasibility. Compared to linear decoders or data-hungry models like recurrent neural networks and transformers, our PSID-based decoder provides a clinically viable and computationally efficient alternative. Despite its simplicity, the decoder generalized over a range of tasks and contexts. For example, the decoder trained on a 2D center-out task enabled successful control in obstacle and respawn tasks, despite these involving obstacle configurations and respawn dynamics that were not present during training. Moreover, performance on novel targets (not included in the training set) was comparable to—or even better than—that on pretrained targets. Furthermore, the decoder successfully transferred to different environments and visual perspectives: although trained on a sphere moving on a grid, the decoder supported online decoding in a visually distinct forest environment and functioned with a monkey avatar and in first-person perspective. Such generalization suggests that the decoder captured abstract, behaviorally relevant neural dynamics rather than relying on low-level visual or contextual cues.

Importantly, we observed a persistent performance gap between offline and online decoding—despite using identical decoders and neural data—which strongly implicates neural adaptation as a driver of improved online control. This interpretation is supported by our within-session trend analysis, where monkeys (particularly Monkey 1) showed progressive within-session performance improvements under fixed-decoder conditions. These adaptations likely reflect closed-loop learning mechanisms, where real-time feedback facilitated dynamic adjustment of neural output. Offline simulations, even when using full channel sets, could not replicate these gains, emphasizing the necessity of real-time interaction for capturing the full capabilities of BCI systems.

Another important comparison concerns decoding performance during passive fixation relative to during natural movements. In a previous study from our lab(*32*), we used a similar decoder in a center-out task on a 2D screen and found that performance was higher when the monkey made overt movements compared to a passive condition (average across Monkey 1 and 2: 63% and 53%, respectively). However, that study used a channels selection paradigm and online ReFIT recalibration. In the present study, we used a more complex task without recalibration and trained the decoder on all spiking channels. Despite these additional challenges, success rates in our passive condition were comparable to—or even exceeded—those in the movement-assisted experiment by our group(*32*). This underscores the robustness of our decoder and shows that high-performance BCI control is possible without movement or proprioception.

Several biological and technical design choices likely contributed to this success. Unlike most iBCI studies that rely solely on M1 and PMd, we simultaneously recorded from M1, PMd, and PMv. A parallel study(*32*) showed that PMv activity alone can support online cursor control at performance levels comparable to M1 or PMd. Our offline analysis showed that premotor activity from PMv and PMd carried rich, behaviorally relevant information for BCI decoding. Using matched numbers of high-weight channels we found that PMv and PMd consistently outperformed M1 across tasks and monkeys. The combination of PMv+PMd often matched or exceeded the “All” condition (balanced PMv/PMd/M1 input), suggesting a synergistic premotor contribution and underscoring the limited independent contribution of M1 if premotor signals are available. These findings align with our decoder weight analysis, where most decoding weight was assigned to PMv and PMd (e.g., 60% PMv in Monkey 1), and further validate PMv as a viable and underutilized BCI signal source.

Additionally, our 3D VR environment with stereoscopic vision likely improved the realistic visual feedback to the animals, improving the closed-loop learning. We achieved depth perception by means of liquid crystal shutters operating at 60 Hz and synchronized to the refresh rate of the monitor, a setup in which monkeys achieve excellent performance in a 3D-structure categorization task(*33–35*) suggesting optimal stereo vision. In addition, the VR environment created in Unity contained many other depth cues (perspective, shading, texture, relative motion) similar to real-world situations.

Altogether, our iBCI system demonstrates several critical capabilities that make it highly suitable for assisting paralyzed patients in real-life situations. It enables fast, precise control and continuous navigation, which are essential for tasks such as wheelchair control. The system can seamlessly adjust to changes in target locations and successfully perform obstacle avoidance, all of which are necessary for navigating in complex, cluttered environments such as a house, while also responding quickly to changes in the patient’s intentions. Our BCI was also capable of generalizing across different environments, transitioning smoothly from a basic environment to a complex virtual environment without additional training. This adaptability, combined with a short straightforward Passive Fixation phase, enhances its practicality for home use. Additionally, our BCI’s performance was unaffected by muscle activity, ensuring reliable control even in patients with residual movement. The dominant role of PMv and PMd in decoding, coupled with clear evidence of online neural adaptation, suggests a powerful cortical substrate for intuitive, closed-loop control. These features suggest that our iBCI could substantially improve the quality of life for paralyzed patients by enabling them to perform daily tasks with greater independence.

## Materials and Methods

### Surgery and recording procedures

Three male rhesus monkeys (Macaca mulatta, 7-9kg) were implanted with a titanium headpost that was fixed to the skull with titanium screws and dental acrylic. After training the monkey in a fixation task, we implanted three 96-channel Utah arrays with an electrode length of 1 or 1.5mm and an electrode spacing of 400µm (4×4mm, Blackrock Neurotech, UT, USA), guided by stereotactic coordinates and anatomical landmarks. The arrays were inserted with a pneumatic inserter (Blackrock Neurotech, UT, USA) with a pressure of 1.034bar and an implantation depth of 1mm. During all surgical procedures, the monkey was kept under propofol anesthesia (10mg/kg/h) and strict aseptic conditions. Postoperative anatomical scans (Siemens 3T scanner, 0.6mm resolution) verified the positions of the Utah arrays in dorsal premotor area F2, ventral premotor area F5c, and the hand/arm region of the primary motor cortex. All surgical and experimental procedures were approved by the ethical committee on animal experiments of the KU Leuven and performed according to the National Institute of Health’s Guide for the Care and Use of Laboratory Animals and the EU Directive 2010/63/EU.

During a recording session, neural data were collected with digital Cereplex M headstages (Blackrock Neurotech, UT, USA) that were connected to two digital neural processors. The data were subsequently sent to the Cerebus data acquisition system that recorded the signal of 256 channels, thus allowing simultaneous recording of the three brain areas (one bank of 32 electrodes in PMd was not recorded). The signal was high pass filtered (750Hz) and sampled at 30 kHz. Each recording day, the thresholds to detect spikes were set manually below the noise to capture the neural activity of individual neurons. Note that one channel could contain the signal of one or multiple neurons.

### Experimental setup

The monkey sat in a chair with its head fixed and arms restrained in front of a Viewpixx 3D screen (VPixx Technologies Inc., Saint-Bruno, Canada, 1920×1080 pixels with a refresh rate of 120 Hz) in a dark room (Fig.4). On the screen, pairs of images for left and right eye with slight angular disparities were presented alternately at 120 Hz. The monkey wore shutter glasses with liquid crystal lenses that opened and closed when voltage was applied, which was perfectly synchronized with the screen at a frequency of 60Hz for each eye to achieve stereoscopic vision. A similar setup was used in detailed behavioral and electrophysiological testing of 3D vision based on disparity in macaque monkeys(*33–35*). An infrared camera monitored the monkey’s eye movements with a sampling rate of 500 Hz (EyeLink 500, SR Research, Ontario, Canada). Prior to each recording session, the eye tracking system was calibrated by requiring the animals to fixate targets presented on the 3D screen.

**Fig. 4.**
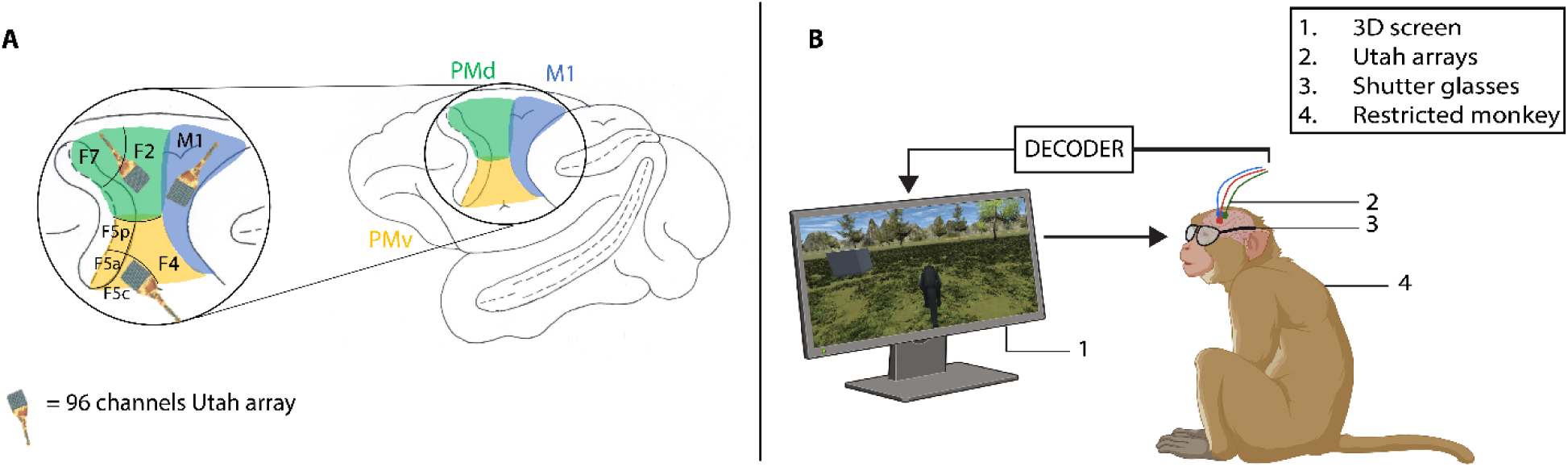
Implantation site and experimental setup. **A** Implantation site of three 96-channels Utah arrays: one in PMd, PMv and M1. **B** Experimental setup of closed-loop BCI: head-fixed monkey sitting in a chair with restrained arms. A 3D screen in front of the monkey displayed the task. The Monkey wore shutter glasses synchronized with the 3D screen, to allow the perception of depth.

### Experimental design

Each session consisted of a Passive Fixation phase, a Decoder Training phase and an Online Decoding phase. The same decoder was used across tasks unless the task geometry or perspective changed, in which case a new decoder was trained. The main goal of each experiment was to move an entity (a sphere or a monkey avatar) from a starting point to a target that pseudorandomly appeared in the 3D environment.

#### Experimental phases

In the Passive Fixation phase, the monkey passively observed Unity’s task execution using its AI-driven navigation system. This phase employed three strategically placed targets—left, straight ahead, and right—to cover the main directions in the 3D environment. For each target, we typically collected 30 trials while recording neural activity in three motor areas (M1, PMv and PMd) and the velocities of the moving entity. In the Decoder Training phase (typical duration of a few sec to 1 min), we trained a model using the neural activity and the entity’s velocities that were recorded during the Passive Fixation phase. In the Online Decoding phase, the monkey controlled the entity using its decoded neural activity. The tasks developed in Unity used computed velocities from the decoding algorithm as inputs, enabling real-time control of the entity’s movement. The online decoding formed a closed-loop Brain-Computer Interface (BCI) as the monkey observed the decoded velocities in real-time on the 3D screen.

#### Tasks

The first task was a Center-out task, featuring a sphere in a 3D environment that moved on a two-dimensional (2D) plane towards the target (a white cube, Fig.5A). During the Passive Fixation phase, the sphere moved from a fixed starting point in a straight line to one of three targets (far left, straight, and far right) that appeared pseudorandomly. The eyes were monitored to ensure fixation on the part of the screen where the sphere moved towards the target. For each target, 30 correct trials were recorded. During the Online Decoding phase, the monkey needed to reach five possible target positions (Fig. S2) that appeared pseudorandomly by controlling the sphere movements through the decoding algorithm.

**Fig. 5.**
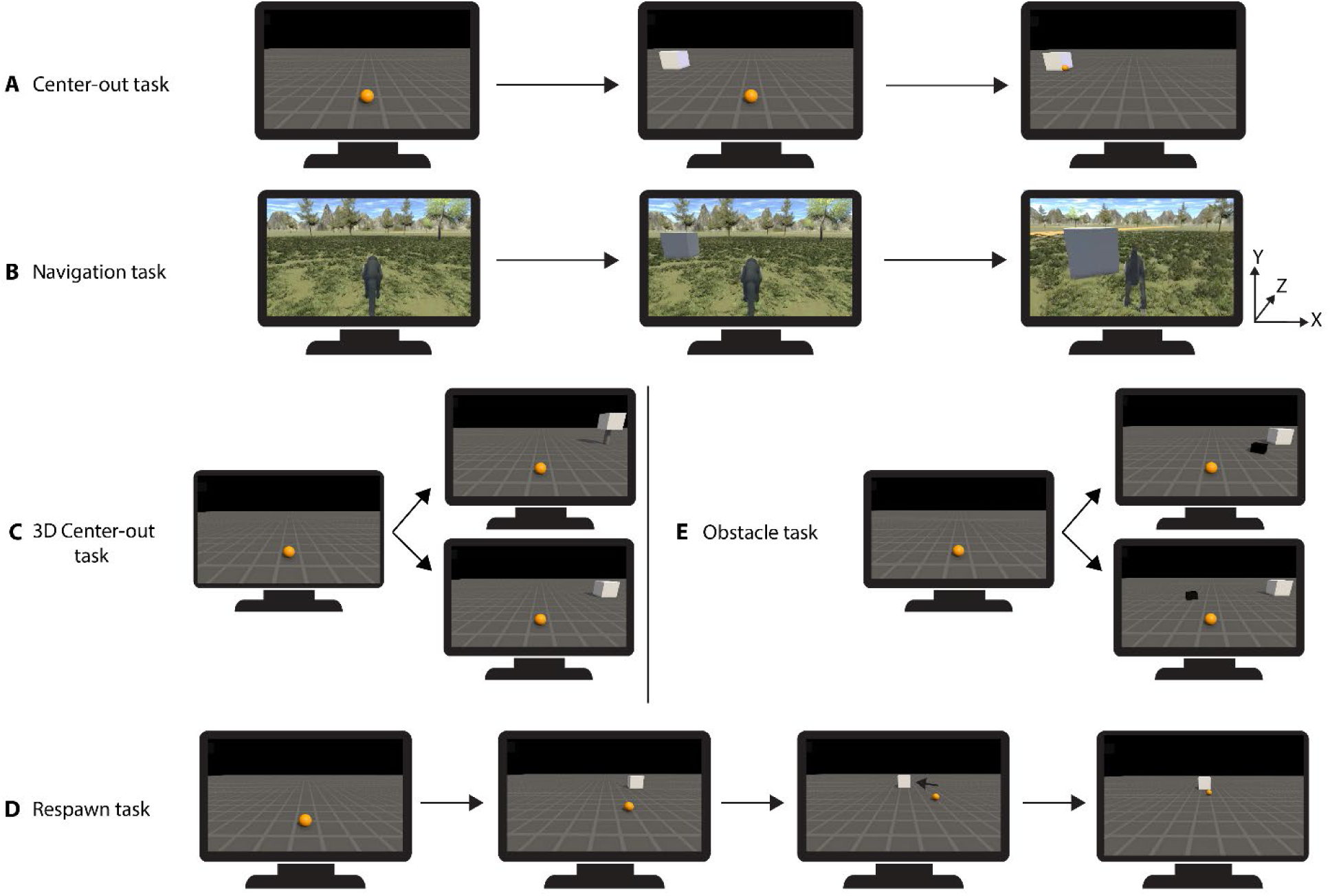
Experimental tasks. **A-B** Two main tasks: temporal sequence of the Center-out task (A) and Continuous Navigation task (B) in xyz-space. **C-D** Three supplementary tasks: 3D Center-out task with two possibilities, upper target and lower target (C). **D** Respawn task in chronological order, with the respawning of the target to a different location during the decoding trial. **E** Obstacle task with two possibilities during the decoding trial, above with obstacle on the path of the target and below with an obstacle off the path of the target.

In the Continuous Navigation task, the starting position at the beginning of each trial was the end position of the previous trial. In the Passive Fixation phase, the monkey observed a sphere performing continuous movements on a 2D grid. The sphere followed a curved trajectory towards the target, and the camera dynamically tracked the sphere on the grid (Supplementary Movie S7). The Center-out task and the Continuous Navigation task differed with respect to the camera movement. In the Center-out task, the camera remained fixed, the sphere moved in the z-(depth) and x-direction (width) and the targets remained at a constant distance from the camera, causing the sphere to appear smaller on the screen as it moved toward the targets. In contrast, in the Continuous Navigation task, the camera dynamically tracked the sphere on the grid. As the sphere approached the targets, their size increased on the screen. Similar to the Center-out task, three targets were used during the Passive Fixation phase, and 30 correct trials of each target were recorded. While the Passive Fixation phase was performed with a sphere moving on a plane with dynamic camera tracking, the Online Decoding phase used a monkey avatar navigating in a 3D forest environment (Fig.5B). When the avatar rotated, the camera rotated correspondingly, revealing different sections of the 3D forest. During the Online Decoding phase, the monkey was required to navigate the avatar monkey to five different target positions in the forest environment. During this phase, we recorded 100 trials per session, for both main tasks.

Monkey 2 and 3 performed three additional tasks that were based on the main Center-out task. In the first additional task, the sphere was moved in the *x* (width), *y* (height), and *z* (depth) directions (3D Center-out task). This task used ten different targets, five of which were situated on the plane at identical positions to those in the first Center-out task and the remaining five targets shared the same *xz*-coordinates but had positive *y*-coordinates (i.e. located higher than the five targets on the plane, balancing on thin cylinders, Fig.5C). In the Passive Fixation phase of the 3D Center-out task, six targets were shown; three on the ground and three elevated, and for each target we recorded fifteen correct trials. During the Online Decoding phase, the animal moved the sphere in three dimensions to reach each of the ten possible targets that appeared pseudorandomly.

For the last two tasks, the Passive Fixation phase was identical to the Center-out task. During the Online Decoding phase of the Respawn task, the target could respawn (i.e. disappear and reappear at another location) to one of its adjacent target positions after the sphere crossed a specific point in the space (*z*-coordinate equal to two, Fig.5D). Thirteen target patterns were possible and chosen pseudorandomly: no respawning of the five target positions, and respawning of each target in two possible directions (to the left or to the right) or in one possible direction for the most left and most right target. In the Obstacle task, an obstacle was added during the Online Decoding phase (Fig.5E). The obstacle appeared pseudorandomly on or off the path to the target. The obstacle was smaller than the target, situated in the middle of the trajectory to the target and oriented in the direction of the specific target, so that the sphere had to deviate from a straight trajectory in on-path trials. During the Online Decoding phase of the three additional tasks, 120 trials were recorded per sessions, balanced across the different task conditions explained above. The dimensions of all tasks can be found in Fig. S2.

#### Experimental protocol

In the Center-out tasks, the sphere appeared at the starting position on the *xz*-plane, while for the Continuous Navigation task, the starting position of the sphere/avatar was the end position of the last trial. To start a trial, the monkey had to fixate on the sphere or avatar for a duration of 500ms. Subsequently, a target appeared at one of the predetermined positions. The sphere or avatar had to reach the target within a predefined time period, denoted as maximum trial time (5.5s for the Center-out task and the 3D Center-out task, 8s for the Continuous Navigation task and 6.5s for the Respawn and Obstacle tasks). A trial was successful when the sphere or avatar stayed inside a predefined box around the target (target window) for 500ms. Successful trials were rewarded with a liquid reward. An unsuccessful trial occurred if the target was not reached within the maximum trial time or if the sphere, avatar, or target exited the camera’s field of view. Trials in which the subject failed to maintain fixation at the beginning of the trial were aborted and not further analyzed. Upon completion of a trial in the Center-out tasks (whether successful, unsuccessful, or aborted), both the sphere and the target disappeared from the screen. If in the Continuous Navigation task the avatar did not reach the target in the predefined time window or if the target went out of the scene, the trial stopped and the avatar remained at its final position, while the target disappeared.

### Unity

#### Software and hardware

The tasks were developed using Unity 2021.3.16f1 and implemented on standard desktop computers. The Unity’s 3D Universal Render Pipeline (URP) template was used as a starting point for developing each task. The subject interacted with the tasks developed in Unity using a custom-built BCI system. Unity was installed on a first computer controlling the 3D screen that the monkey was looking at. On a second computer, the decoding model (in python) was trained to decode velocities in real-time. On a third computer, the central task script (in python) was executed, facilitating the exchange of commands between the three programs. These commands included basic instructions such as configuring trials, starting and stopping trials, and predictions. A Local Area Network (LAN) facilitated communication between the three programs using ZeroMQ over TCP/IP sockets to enable efficient and reliable data exchange in a distributed system.

#### Scene composition

The Center-out task was set up in an environment with a flat plane featuring a grid pattern, providing multiple depth cues in 3D space. The sphere was able to translate in the *x* and *z* directions in the Center-out, Respawn and Obstacle tasks and in the *x*, *y*, and *z* directions for the 3D Center-out task. The targets were white static cubes. In the 3D Center-out task, the elevated targets were located on thin cylinders. In the Continuous Navigation task, the environment was a virtual forest in an open field bordered by mountains (Fig.5B). The navigational entity was now represented by a monkey avatar (a capuchin monkey) obtained from the Unity store(*36*). The dimensions of each scene are shown in Fig. S2.

#### Movement during training

The movement of the sphere and avatar during the Passive Fixation phase of the experiment was controlled by a built-in Unity component, the Nav Mesh Agent. Attached to the sphere/avatar game object, the Nav Mesh Agent operated on a baked Nav Mesh on the navigation plane. The ‘SetDestination’ method was utilized to update the Nav Mesh Agent’s destination, initiating the autonomous movement of the sphere towards the target in a straight trajectory. The motion began with an acceleration, gradually reaching a steady speed of four units per second. As the sphere approached the target position, it decelerated, ensuring that the sphere came to a complete stop just before making contact with the target. For the Continuous Navigation task, the sphere moved along a smooth, curved trajectory towards the target. Our method leveraged Bezier curves to compute a curved path towards the target. The trajectory path was divided into destination points, and the ‘SetDestination’ function guided the avatar through these points, resulting in a realistic and more natural movement pattern. The movement during the 3D Center-out task involved the sphere moving on the plane (to the lower targets) and flying from the ground to the elevated targets. The use of the nav mesh agent was not possible due to its restriction to the surface-only baked nav mesh. Instead, the Unity build-in function ‘transform.Translate’ was used to move the sphere in a straight line, along with the simulation of acceleration and deceleration.

#### BCI-controlled movement

The decoding algorithm produced velocities (*v_x_*, *v_y_*, *v_z_*) that dictated the direction and the speed of the sphere/avatar’s movement. These velocities were transmitted to Unity every 50ms, forming the basis for real-time updates to the sphere/avatar’s position in the virtual environment. The sphere and the avatar were equipped with a rigid body component. The decoding algorithm dynamically adjusted Unity’s rigid body velocity, translating navigational intent into realistic sphere and avatar movements. The sphere/avatar were only responsive to the decoding algorithm’s output during the interval from target onset to target acquisition. The sphere was limited to translation in the *x*, *y* (for the 3D task), and *z* directions. In the Continuous Navigation task, the avatar could translate in the *x* and *z* directions while dynamically rotating based on the velocities. From the *x* and *z* velocities, we computed the rotation angle and forward speed. The forward speed (*v*) was computed as the magnitude of the velocity vector using the formula: 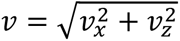. The rotation angle (θ) was calculated using the arctangent function: 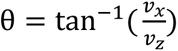. Rotation was achieved using the ‘Quaternion.Lerp’ function applied to the avatar’s transform, ensuring a smooth rotation that corresponded to the avatar’s actual movement.

#### Camera view

A dual-camera setup replicated the visual perspectives of the left and right eyes to create binocular disparities. These perspectives were sequentially displayed on the 3D screen, simulating a stereoscopic or 3D view. The camera projection was configured for a perspective view, and the field of view for each camera was set to 60 degrees. For the Center-out task, the camera positions remained fixed. In contrast, during the Continuous Navigation task, the cameras dynamically tracked the avatar in the 3D environment. The tracking was achieved using the command ‘transform.LookAt()’, enabling the camera to smoothly translate and rotate in sync with the avatar’s movements. Throughout the task, it was essential to verify that the sphere (for the Center-out tasks) or the target (for the Continuous Navigation task) remained within the camera’s field of view at all times. To achieve this, the Unity function ‘GeometryUtility.TestPlanesAABB(camera_frustum, collider.bounds)’ was employed. This function assessed whether the collider bounds of the object (sphere or target) remained within the 3D space visible to the camera (camera fustrum). The camera frustum was computed with the Unity function ‘GeometryUtility.CalculateFrustumPlanes’. This evaluation was conducted for both the left and right eye camera views.

#### Reaching the target

The target was reached by the sphere/avatar when it stayed within a predefined box around the target (target window) for 500 milliseconds. Since both the target and the controlled sphere are 3D physical objects that could not intersect, we defined the target window based on the distance from the target’s edge to the window boundary, which was 1.35 units, with the sphere’s diameter being 0.5 units. The larger window also accounts for the requirement that the sphere must remain inside it for 500ms, allowing for minor movement during this time. The dimension of the target window for the different tasks are shown in Fig. S2. During the Respawn task, the target’s position and rotation were changed to the new position when the sphere crossed the grid coordinate equal to two. For the Obstacle task, an additional cube was added to the scene around which the sphere needed to navigate to reach the target.

#### Avatar animation

In the Continuous Navigation task, the monkey avatar was equipped with walking, running, and idle animations. The monkey avatar adopted an idle animation, resting on both its arms and legs without any motion, when its velocity was at zero. If the avatar’s velocity was higher than zero and below two units per second, the walking animation was activated. When the velocity exceeded 2 units/s, the running animation took over. The speed of the running animation was adjusted based on the actual velocity, providing visual feedback that reflected the avatar’s current speed.

#### Additional functions

The Continuous Navigation task contained virtual trees in the forest environment. The avatar could be blocked behind a tree if the subject did not navigate properly around it. To ensure the task focused on reaching the targets and not avoiding trees, a special function made sure that if the avatar stayed blocked behind a tree for the entire trial, its starting position was reset to the center of the forest for the next trial. This reset was initiated by checking the avatar’s velocity (near zero for over four seconds). The environment in the Continuous Navigation task, whether the grid or the forest, was not infinite. If the avatar was close to the environment’s edge, it was reset to the center at the start of the next trial. This was controlled by fixed boundaries to define the environment limits.

### Decoding algorithm

The decoding algorithm translated neural activity of PMv, PMd and M1 into the movement of the sphere/avatar in real-time. Initial training of the decoder relied on labeled training data, employing a nonlinear extension of the Preferential Subspace Identification (PSID) framework(*37*), which we adapted for real-time, closed-loop BCI. While PSID was originally developed for offline analysis, we extended it by adding a non-linear regression stage to decode continuous 3D velocities. Once the decoding algorithm was trained, it was able to predict real-time velocities based on real-time brain signals. This combination of nonlinear decoding and real-time state estimation enables fast, interpretable, and task-generalizable BCI control.

#### Electrode selection

With our data acquisition system, we could record from 256 electrodes simultaneously. Before each experiment, electrode signals were visually inspected, and if no spikes were detected, the spike threshold was set below a predefined value. During the training of the decoding algorithm, electrodes with thresholds below the predefined value were excluded from the model.

#### Training

The training of the decoding algorithm involved a supervised method using labeled data from the Passive Fixation phase. The data included spike rates of each selected electrode for each 50ms time bin, along with the corresponding velocities (*v*_*x*_, *v*_*y*_, *v*_*z*_) and positions of the sphere/avatar at that timestamp. The PSID method was employed to train the decoding algorithm. The state of the brain was considered as a high-dimensional latent variable at each time point. A dynamic linear state space model was used to characterize both the neural activity and the behavior. Since we only wanted to consider neural dynamics that were related to the behavior of interest, we computed the behaviorally relevant latent variables (Fig. S6A) together with the corresponding model parameters in this Decoder Training phase using two parameters: the number of extracted states *n*_1_ = 6 and the projection horizon *i* = 5 (*37*).

After extracting latent variables from the neural signal, the next step involved deriving velocities from these latent variables using regression (Fig. S6B). Due to the inherent non-linear relationship between the latent variables and velocities, a kernel approximation was employed as a preprocessing step. Using the Nystroem kernel approximation with the Radial Basis Function (RBF), the data underwent a projection into a higher-dimensional space, enhancing the capturing of non-linear patterns. The Nystroem method approximates the kernel matrix using a smaller subset of data points, reducing the computation time. The chosen parameters of the Nystroem kernel approximation were: γ = 0.34 that influenced the smoothness of the decision boundary of the Gaussian kernel, *n_components_* = 700 was the number of data points used in the approximation and *random_state* = 42 was used for reproducibility, ensuring that the same random subset of data points was selected in each run. Next, a linear regression was applied to predict the velocities. The linear regression process identified the optimal-fitting flat hyperplane that minimized the difference between predicted and actual velocities. More specifically, a ridge regression was used, which introduced a regularization term to avoid overfitting the model and prevented potential multicollinearity (high correlation) in the data. The strength of the regularization was controlled by the hyperparameter α = 0.1. To ensure that the model learned the relationship between latent variables and velocities based on the provided ground truth, both the kernel approximation and the linear regression were trained using the labeled training data. The different parameters used for the training of the decoder were computed using hyperparameter optimization on a subset of the training data in a preliminary pilot study.

#### Online decoding

To predict velocities from neural activity in real-time, a recursive Kalman filter was used in combination with the regression step. The recursive Kalman filter estimated and updated the latent states of the system based on the observed brain signals in a sequential and iterative manner. The filter processed each new observation, in this case the neural activity in a 50ms time bin, and refined its estimate of the latent states based on the current and past information. The principal advantages of using a recursive Kalman filter are the sequential updating of the latent states, the ability to handle noisy observations and to adapt to changing dynamics over time.

### Statistical Analysis

#### Success rate and chance level

The success rate was defined as the ratio of successful trials to the total number of trials for each session. To assess the significance of differences in success rates across various groups, we calculated Mann-Whitney U tests. The groups compared were the upper and lower targets for the 3D Center-out task, the respawn and no respawn trials for the Respawn task and the on-path and off-path obstacle trials for the Obstacle task (all taken across all sessions). To estimate the chance level of each task, we conducted a permutation test. For each 10,000 permutations, the target labels were shuffled and a test statistic—in this case, the success rate across trials— was computed. By comparing the observed statistic (success rate of the monkey across all trials and sessions) to the distribution of the permuted statistics, we could quantify the likelihood of observing such performance by chance. To determine if the observed success rate was significantly different from what could occur by random chance, we computed the p-value by determining the proportion of permuted test statistics that were as, or more, extreme than the observed success rate.

#### Offline decoding analysis per cortical region

To assess the contribution of the different cortical areas to decoding performance, we performed an offline analysis using data from the same sessions as the Online decoding experiments. For each session, we selected the subset of electrodes with the highest cumulative decoder weights, as defined by their absolute contributions to the latent variables in the trained online decoder. To ensure comparability across regions, we used the same number of channels for each area, based on the minimum available channel count across M1, PMd, and PMv (Monkey 1: 27-35, Monkey 3: 41-61 channels). Decoding performance was then evaluated using neural signals from each area individually (M1, PMd, PMv), from pairwise combinations of areas (PMv+PMd, M1+PMd, M1+PMv), and from a balanced selection across all three regions (“All”). These configurations were compared to two references: a decoder using all channels offline (“Online simulation”), and the actual performance achieved in the closed-loop online condition (“Online decoding”). Decoding success was quantified and normalized either to the “All” condition, the “Online simulation”, or the “Online decoding” performance, allowing us to assess the relative area contributions, and evaluate the impact of closed-loop adaptation. Statistical comparisons between conditions were performed using Wilcoxon signed-rank tests (two-sided), with a significance threshold of *p* < 0.05.

In addition, we quantified the relative contribution of each brain area to the trained decoder by summing the absolute decoder weights on the latent variables for each electrode. We computed the percentage of total weight contributed by M1, PMd, and PMv separately, based on the decoder trained on all channels. To focus on the most informative electrodes, we applied a 50% cumulative weight threshold: we included electrodes in descending order of their absolute weight until 50% of the total decoder weight was accounted for. This analysis allowed us to assess how strongly each cortical area influenced the decoder, independent of channel count.

Finally, to illustrate the differences between offline and online decoding performance, we integrated these results with our within-session adaptation analysis (see next section), which quantifies the extent to which monkeys improved performance under fixed-decoder conditions within a session.

#### Within-Session Adaptation Analysis

To assess whether monkeys adapted their neural activity to improve control within a session and in a given task under fixed-decoder conditions, we analyzed trial-by-trial task performance. For each session, a smoothed success rate was computed using a moving average (window = seven trials, step size = 1). We then fit a linear regression to the smoothed success rate across all trials in the session. Sessions were classified based on the slope and significance of the fit: Improvement (positive slope, p < 0.05), Decline (negative slope, p < 0.05) or Constant (p ≥ 0.05). This analysis was performed for each monkey–task combination, and results were then summarized both per monkey and pooled across monkeys for each task.

#### Reaction time

We computed the reaction time in the Respawn task as the time interval between the change in target position and the detected point of deviation in the sphere’s trajectory, which corresponds to the monkey’s response to the target change. We started by smoothing the trajectory using a low-pass Butterworth filter and a moving average. Next, we computed the tangents along the trajectory to determine the movement direction at each time point following the target jump time. The reaction point was defined as the first noticeable deviation in movement direction after the target jump, identified using a predefined angular threshold. Importantly, we accounted for the ∼100ms delay between the neural intention to change direction and the resulting visible movement of the sphere on the screen (because of delays introduced by neural signal binning, software communication, and visual rendering on the display), by subtracting it from the reaction time.

#### Time to target

We calculated the time it took the monkey to move the sphere/avatar to the target. This time to target measurement started from the moment the sphere/avatar was allowed to move and ended when the trial was successful, defined as staying for 500ms within the target window. We compared the time to target during the Passive Fixation phase, where unity was in control, and the Online Decoding phase, where the monkey was in control.

#### Influence of sEMG on decoded velocity

To investigate whether arm and hand movement influenced the decoded velocity during the Training and Online Decoding phase, we recorded sEMG from the biceps (biceps brachii) and the thumb (abductor pollicis brevis) of Monkeys 1 and 2, using two dry-adhesive electrodes, along with a ground electrode placed next to the biceps electrode. The raw sEMG was preprocessed by applying a high-pass Butterworth filter (order = 4, cut-off frequency = 30Hz), a 50Hz notch filter, and a low-pass Butterworth filter (order = 4, cut-off frequency = 500Hz). The signal was rectified, and its envelope was obtained using a Butterworth low-pass filter (order = 4, cut-off frequency = 20Hz)(*38*). For every trial, the sEMG signal was segmented into 50ms bins to align with the decoded velocity, averaged over each bin, after which all trials of a session were concatenated.

We assessed the relationship between sEMG and decoded velocities using Spearman correlation analysis. We computed the mean correlation across all sessions and the combined p-value (Fisher’s method) for the different tasks and for the different velocity components (v_x_, v_y_, v_z_, v = magnitude of velocity). To account for potential time delays, cross-correlation analysis was conducted with lags ranging from −100 to +100ms (50ms increments). A positive lag indicates that the sEMG follows the decoded velocity, while a negative lag suggests that the sEMG precedes the decoded velocity, considering typical delays from motor cortex to sEMG (50-150ms(*39*)) and to decoded velocity (50-150ms). Additionally, we performed linear and non-linear regression (polynomial, degree = 2) analyses to evaluate how well sEMG predicted the decoded velocities by computing the *R^2^* and the MSE values, together with the combined p-value.

#### Influence of eye-movements on decoded velocity

During the Training and Online Decoding phase, the monkey watched a screen displaying tasks while its eye movements were monitored continuously. Except for the Continuous Navigation task, the monkey had to maintain its gaze within a large electronically defined window that contained both the initial position of the sphere and the target. For the Continuous Navigation task, the eye-window matched the screen size, because the camera view rotated with the sphere/avatar. To estimate the influence of eye movements on the decoded velocities, we computed the Spearman correlation between the eye position (*x* and *y*-component) and the decoded velocity components (*v_x_*, *v_y_*, *v_z_*, *v* = magnitude of velocity). The eye movement data were segmented into 50ms bins, and the average value was taken as a feature. The correlation coefficients were computed similar to those between the sEMG and the decoded velocity.

## Acknowledgments

We thank Stijn Verstraeten, Marc De Paep, Wouter Depuydt, Inez Puttemans, and Christophe Ulens for technical assistance. We thank Astrid Hermans and Sara De Pril for administrative support.

## Author contributions

Conceptualization: OS, SDS, TD, JGR, PJ

Methodology: OS, SDS, JGR, TD

Investigation: OS, SDS

Visualization: OS Supervision: PJ

Writing—original draft: OS, SDS

Writing—review & editing: OS, SDS, PJ

## Data and materials availability

All processed data and scripts will be made available on the Dryad server.

## Supplementary Materials

### Supplementary Results: Task-Specific time to target Comparisons

**I**n this supplementary analysis, we analyzed the time to target across task-relevant conditions using two-sided Mann-Whitney U tests. In the 3D Center-out task, Monkey 2 showed no significant difference in time to target between upper and lower targets (U = 128357, p = 0.16, N = 988 trials), whereas Monkey 3 required significantly more time to reach the upper targets (U = 58986, p = 1.11 × 10^-6^, N = 630 trials), highlighting individual differences in adapting to vertical spatial control.

In the Respawn task, both monkeys took significantly longer to reach the target on trials involving a mid-trial target jump (Monkey 2: U=142631, p = 1.16 × 10^-12^, N = 960 trials; Monkey 3: U = 90120, p = 3.24 × 10^-10^, N = 775 trials). This increase—less than 330ms for Monkey 2 and 190ms for Monkey 3—is expected due to the need to redirect toward a new target position.

In the Obstacle task, both monkeys exhibited a modest but significant increase in the time to target when obstacles appeared on-path compared to off-path (Monkey 2: U =32963, p = 0.01, N = 630 trials; Monkey 3: U = 49891, p = 0.01, N = 738 trials). Although these differences were statistically significant, the absolute increase in movement time was relatively small (≤ 250ms), suggesting that the monkeys were able to efficiently navigate around obstacles with only minimal impact on performance.

**Fig. S1.**
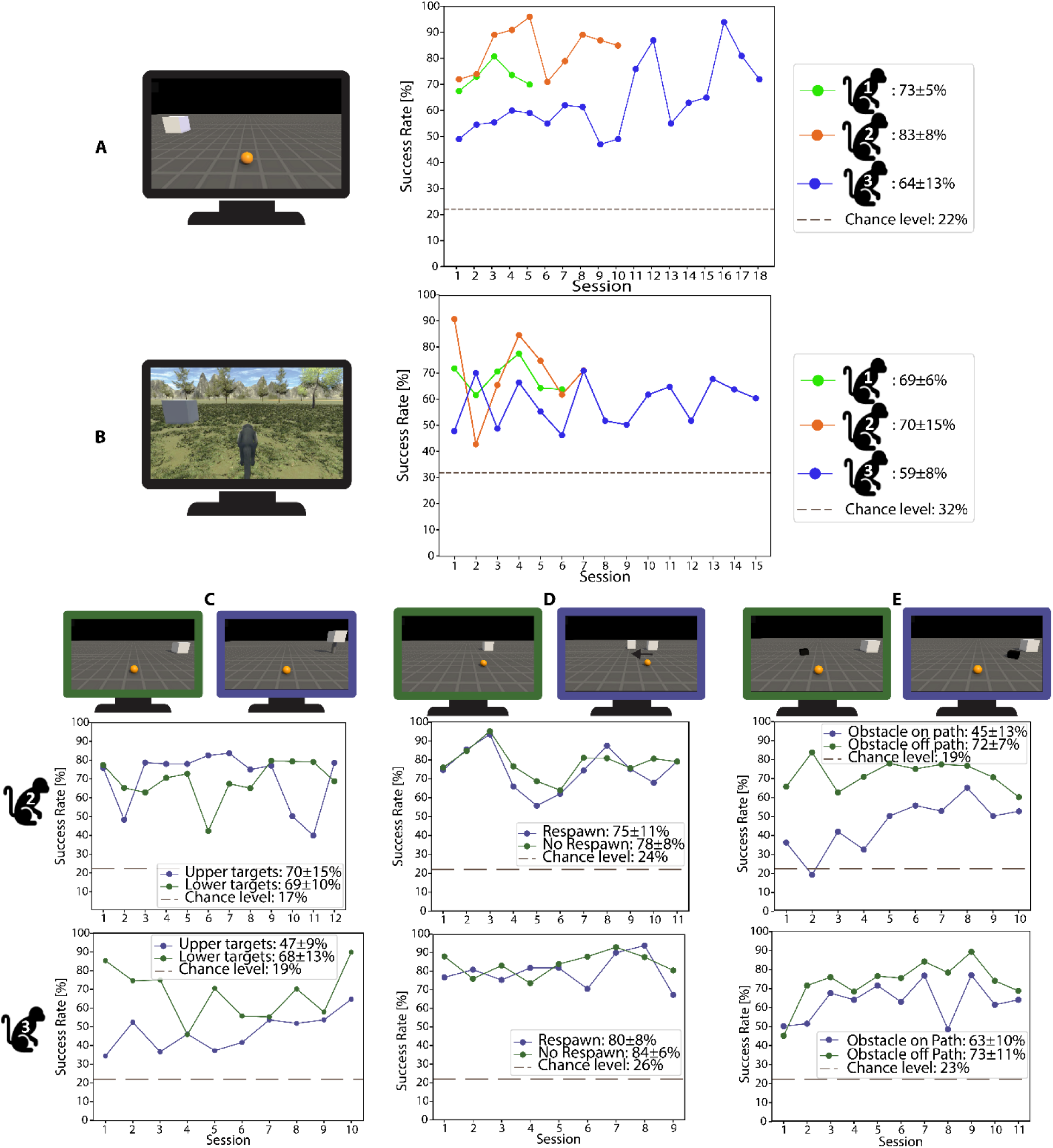
Success rate over time for different tasks. Success rate of each session, together with the estimated chance level and the average success rate ± standard deviation of all sessions. Success rate of Monkey 1, 2 and 3. **A** Center-out task. **B** Continuous Navigation task. **C** 3D Center-out task: upper targets (blue) and lower targets (green). **D** Respawn task: respawn (blue) and no respawn (green). **E** Obstacle task: trials with obstacle on path (blue) and obstacle off path (green).

**Fig. S2.**
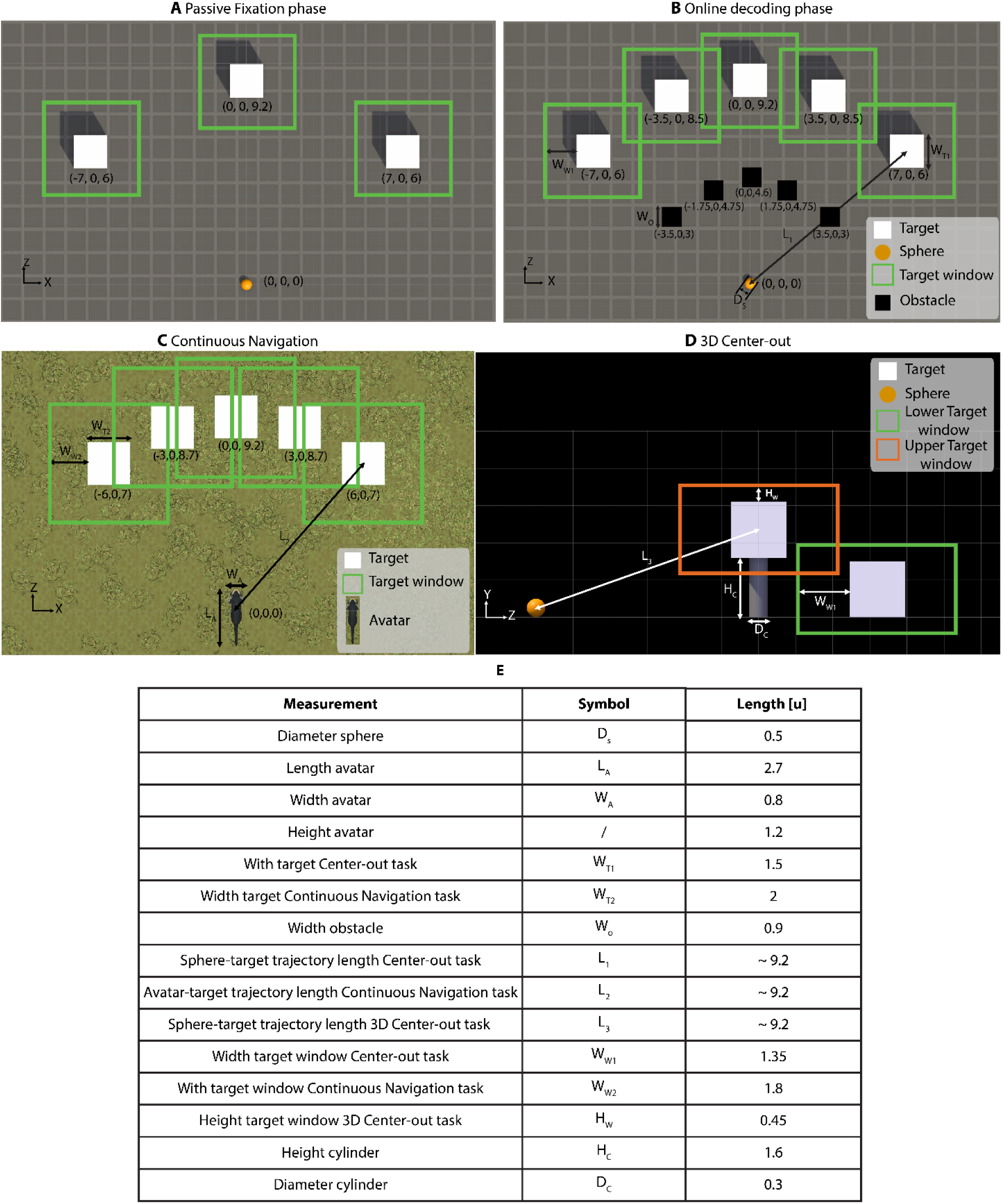
Dimensions, distances and coordinates of main and additional tasks. **A** Passive Fixation phase: only used 3 targets: left, straight and right. **B** Coordinates of sphere, targets and obstacles, together with different dimensions found Table **E**, for Center-out task, Obstacle and Respawn task. **C** Coordinates of sphere and targets, together with different dimensions found in Table **E**, for Continuous Navigation task. **D** Dimensions for 3D Center-out task, values found in Table **E**.

**Fig. S3.**
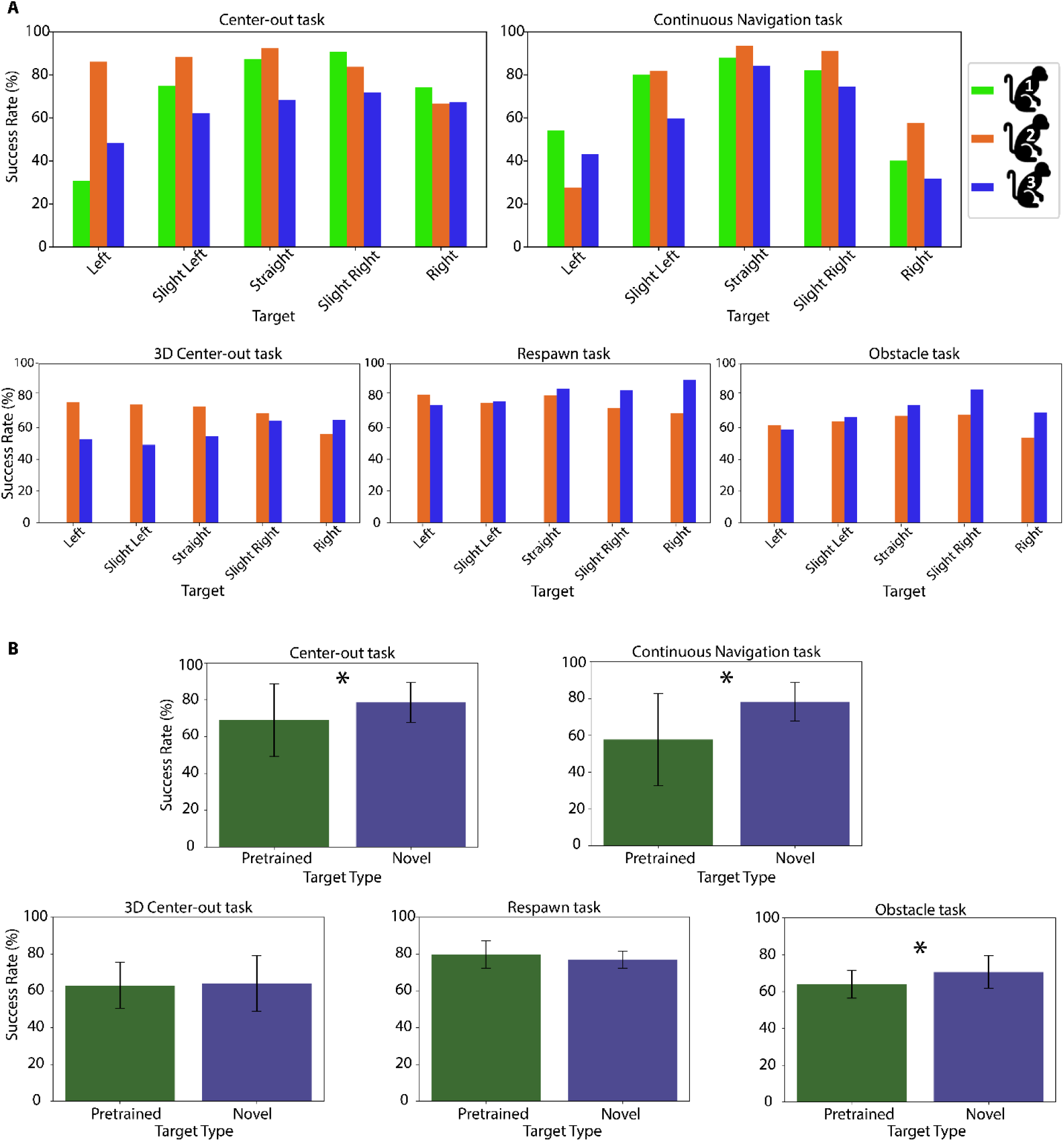
Decoder generalization to novel targets across tasks. **A** Per-target success rate across tasks and monkeys. Each bar shows the mean ± standard deviation success rate per target position for individual monkeys for the different tasks. **B** Comparison of success rates for pretrained targets (left, straight, right) versus novel targets (slight left, slight right). Success rates were calculated per session and averaged across sessions. Statistical comparisons were performed using linear mixed-effects models with monkey identity as a random effect. Performance on novel targets significantly exceeded the performance on pretrained targets for Center-out, Continuous Navigation and Obstacle tasks (p<0.05). In the 3D Center-out and Respawn task there was no significant difference between novel and pretrained targets. The number of sessions (N) for each task is given in Table 1.

**Fig. S4.**
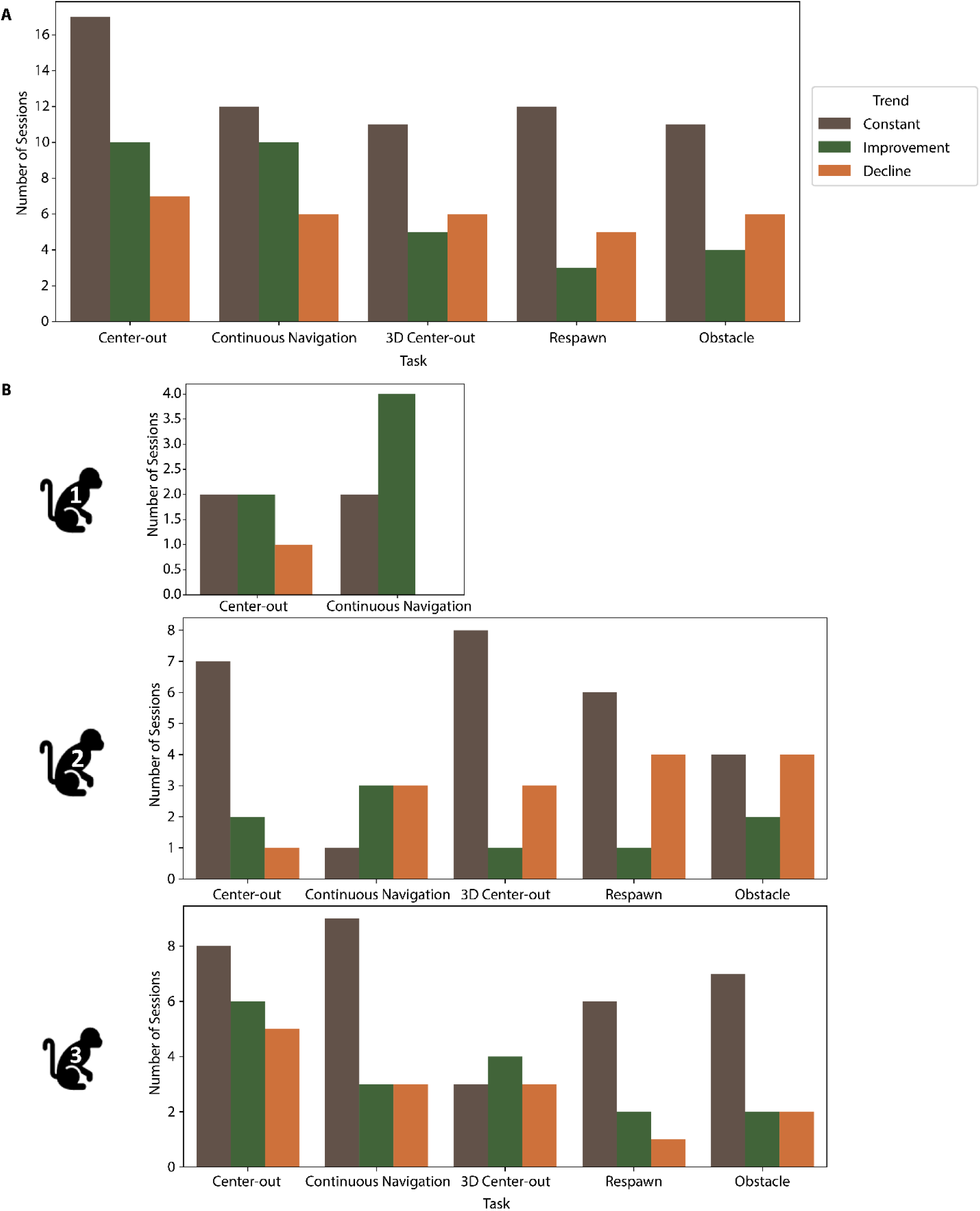
Within-session performance trend analysis across tasks and monkeys. **A** Session-level trend classification (Improvement, Constant, Decline) across all sessions of each task, pooled across monkeys. based on linear fits to smoothed trial-by-trial success rates. Trends were classified by fitting a linear regression to smoothed trial-by-trial success rates and thresholding on slope sign and p-value (Improvement: slope > 0, p < 0.05; Decline: slope < 0, p < 0.05; Constant: p ≥ 0.05). **B** Per-monkey breakdown of trend distributions across tasks. Monkeys differed in adaptation patterns: Monkey 1 showed the highest proportion of improvement and lowest decline, Monkey 2 had more decline sessions, and Monkey 3 showed intermediate behavior. The number of sessions (N) for each task can be found in Table 1.

**Fig. S5.**
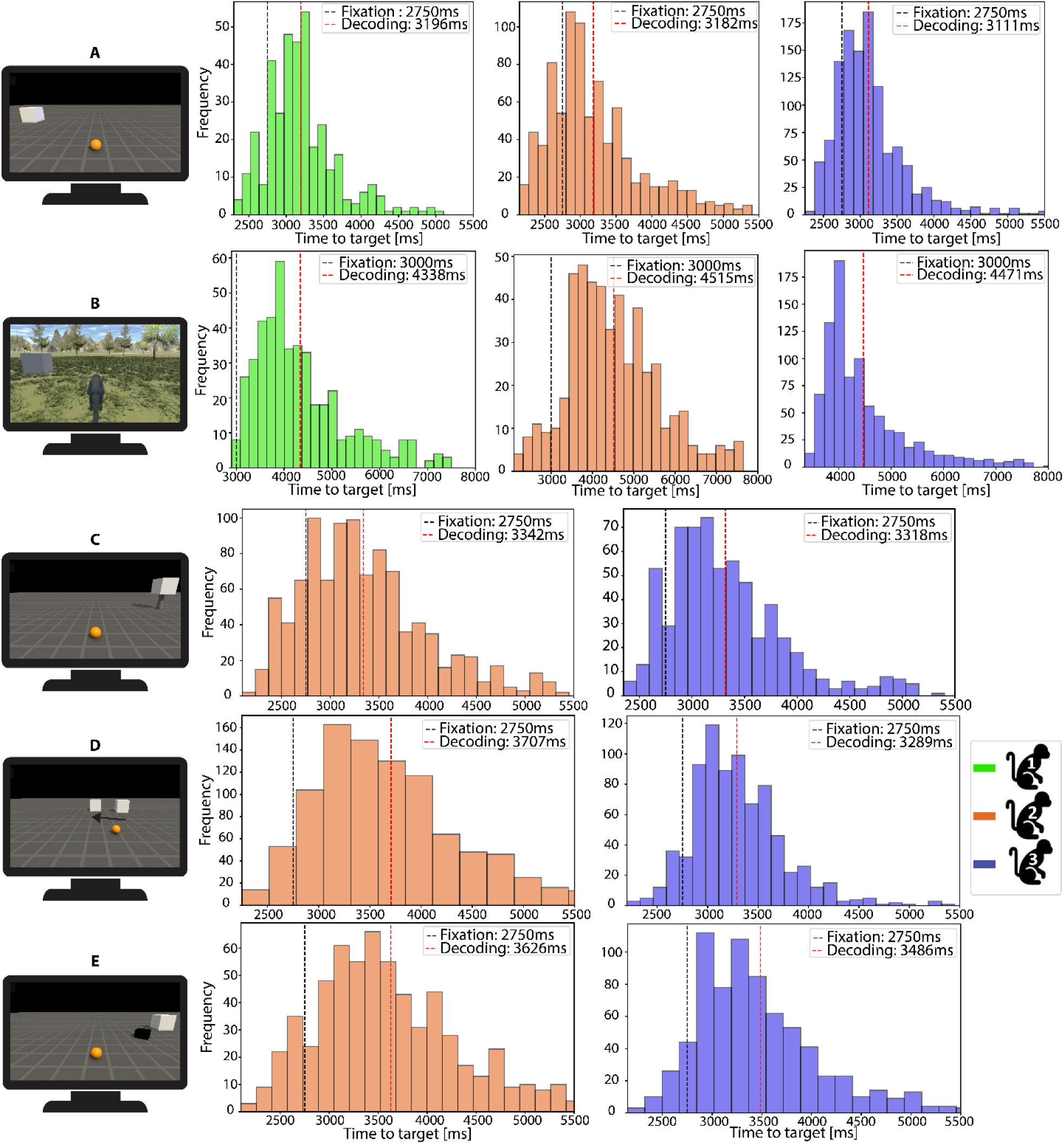
Histogram of time to target for each trial during. Online decoding phase and average time to target for Passive Fixation phase and Online decoding phase, for different tasks and monkey. **A** Center-out task. **B** Continuous Navigation task. **C** 3D Center-out task: time to target on average 21% slower in the Online Decoding phase (compared to the Passive Fixation phase). **D** Respawn task: average time to target in Online Decoding phase was 27% slower compared to the Passive Fixation phase. **E** Obstacle task: time to target was on average 29% slower compared to the Passive Fixation phase. For the Respawn and Obstacle tasks, it is important to note that the Passive Fixation phase did not include changes in target position or the presence of obstacles. The number of sessions (N) for each task is provided in Table 1.

**Fig. S6.**
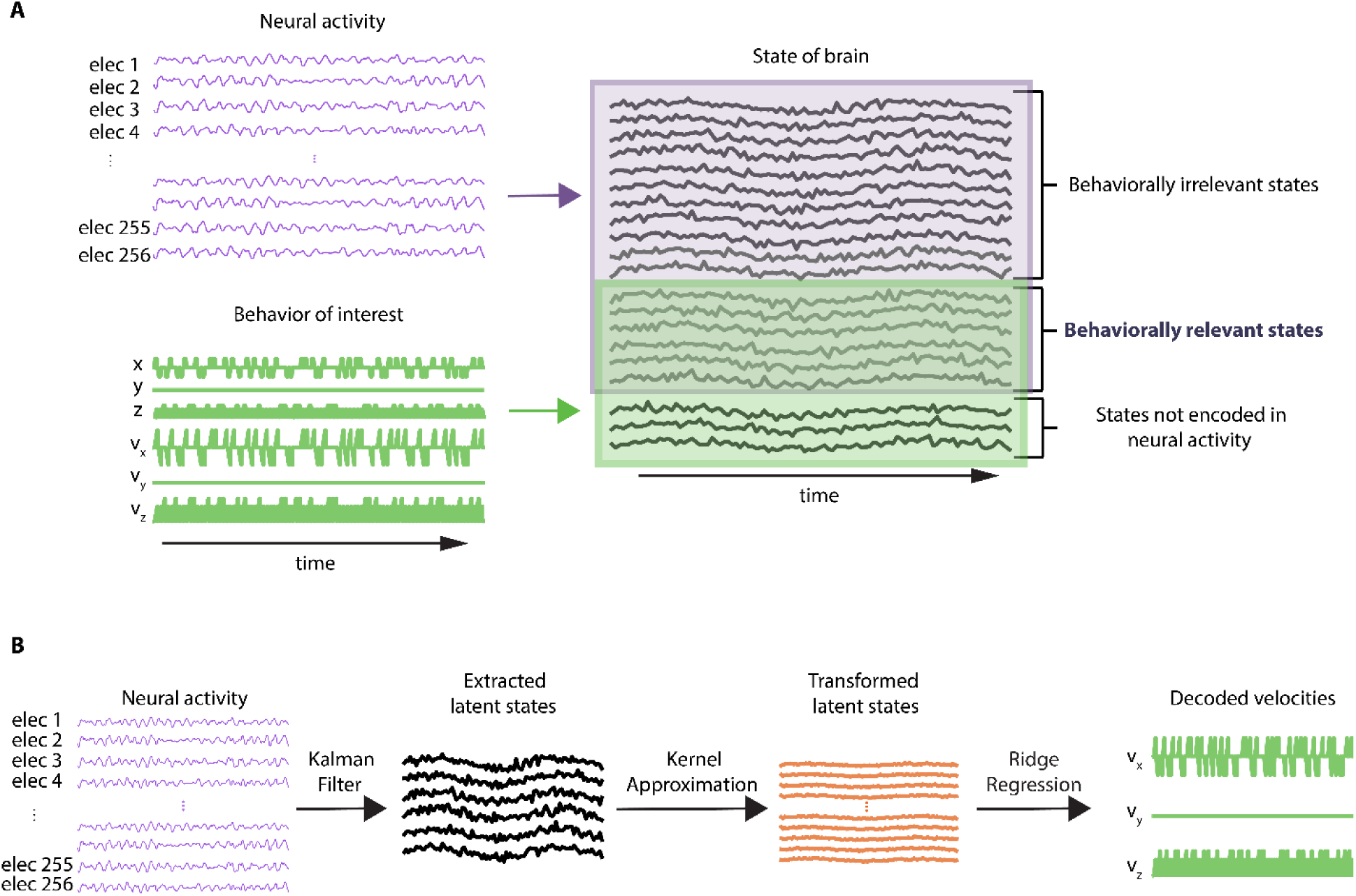
Decoding algorithm scheme. **A** Extraction of behaviorally relevant states. Some dimensions of the latent variable (state of brain) may have driven the behavior of interest (velocity of sphere/avatar, green), some the neural activity (purple) and some both. The dimensions that drive both the neural activity and the behavior of interest are the behaviorally relevant states. Conceptual diagram inspired by (*37*), redrawn to illustrate the separation of latent state dimensions related to behavior and neural activity. **B** Computation of the decoded velocities: starting from the neural activity a Kalman filter was used to extract the behaviorally relevant latent states, then kernel approximation was used to transform these latent states to a higher dimensional space, lastly ridge regression was used to transform the latent states into velocities. These decoded velocities were used to control the sphere/avatar in real-time.

**Table S1.** Statistical comparisons of offline decoding performance across cortical area configurations. Summary of statistical tests comparing decoding success rates across cortical input configurations (see Fig. 2). Tests were performed separately for each monkey and task. For Monkey 1, one-sided exact binomial tests were used due to the limited number of sessions per condition. For Monkey 2, two-sided Wilcoxon signed-rank tests were used to account for repeated sessions. Only statistically significant comparisons are shown (p < 0.05). The number of sessions (N) for each task can be found in Table 1.

| Monkey | Task | Test | Comparison | p-value |
| --- | --- | --- | --- | --- |
| Monkey 1 | Center-out | Binomial (1-sided) | PMd vs M1 | $3.13 \times 10^{-2}$ |
| | | | PMv+PMd vs M1+PMd | $3.13 \times 10^{-2}$ |
| | | | PMd+PMd vs M1+PMv | $3.13 \times 10^{-2}$ |
| Monkey 1 | Continuous navigation | Binomial (1-sided) | PMd vs M1 | $3.13 \times 10^{-2}$ |
| | | | PMv vs M1 | $3.13 \times 10^{-2}$ |
| | | | PMv+PMd vs M1+PMd | $3.13 \times 10^{-2}$ |
| Monkey 2 | Center-out | Wilcoxon (2-sided) | PMd vs M1 | $7.7 \times 10^{-3}$ |
| | | | PMv vs M1 | $4 \times 10^{-6}$ |
| | | | PMv+PMd vs M1+PMd | $1.57 \times 10^{-2}$ |
| | | | PMv+PMd vs M1+PMv | $4 \times 10^{-6}$ |

**Table S2.** Spearman correlation between sEMG and movement velocity components across tasks. This table reports the average Spearman correlation coefficients (mean ± standard deviation) and combined p-values (Fisher’s method) between surface EMG (sEMG) activity and each velocity component (v, vₓ, v_y, v_z) for Monkey 1 and Monkey 2 across multiple tasks. All values reflect pooled session-level statistics for each condition. The number of sessions (N) for each task is provided in Table 1.

| Monkey | Task | Velocity component | Correlation (mean±std) | Combined p-value |
| --- | --- | --- | --- | --- |
| Monkey 1 | Center-out | v | 0.0 ± 0.01 | 0.65 |
| | | v<sub>x</sub> | -0.02 ± 0.04 | 1.95 $\times 10^{-7}$ |
| | | v<sub>z</sub> | -0.01 ± 0.02 | 0.30 |
| | Continuous Navigation | v | -0.01 ± 0.02 | 4.0 $\times 10^{-4}$ |
| | | v<sub>x</sub> | -0.04 ± 0.13 | 2.40 $\times 10^{-127}$ |
| | | v<sub>z</sub> | -0.02 ± 0.03 | 31.69 $\times 10^{-8}$ |
| Monkey 2 | Center-out | v | 0.11 ± 0.13 | 0.0 |
| | | v<sub>x</sub> | -0.13 ± 0.1 | 0.0 |
| | | v<sub>z</sub> | 0.05 ± 0.13 | 2.16 $\times 10^{-198}$ |
| | Continuous Navigation | v | 0.01 ± 0.09 | 1.87 $\times 10^{-57}$ |
| | | v<sub>x</sub> | 0.02 ± 0.12 | 2.28 $\times 10^{-130}$ |
| | | v<sub>z</sub> | 0.0 ± 0.08 | 3.41 $\times 10^{-29}$ |
| | 3D Center-out | v | 0.11 ± 0.11 | 3.25 $\times 10^{-304}$ |
| | | v<sub>x</sub> | -0.2 ± 0.11 | 0.0 |
| | | v<sub>y</sub> | 0.05 ± 0.11 | 1.43 $\times 10^{-196}$ |
| | | v<sub>z</sub> | -0.02 ± 0.1 | 3.23 $\times 10^{-132}$ |
| | Respawn | v | 0.05 ± 0.09 | 9.64 $\times 10^{-146}$ |
| | | v<sub>x</sub> | -0.12 ± 0.13 | 0.0 |
| | | v<sub>z</sub> | -0.06 ± 0.06 | 2.08 $\times 10^{-90}$ |
| | Obstacle | v | 0.09 ± 0.05 | 1.49 $\times 10^{-90}$ |
| | | v<sub>x</sub> | -0.13 ± 0.12 | 4.82 $\times 10^{-293}$ |
| | | v<sub>z</sub> | -0.04 ± 0.09 | 3.37 $\times 10^{-70}$ |

**Table S3.** Regression performance metrics between sEMG and movement velocity components. This table summarizes the results of both linear and non-linear regression analyses relating sEMG signals to different velocity components (v, vₓ, v_y, v_z) for Monkey 1 and Monkey 2 across multiple tasks. Reported metrics include the median R² with interquartile range (IQR), median MSE with IQR, and the combined p-value (computed using Fisher’s method) across sessions. Results are shown separately for linear and non-linear models. The number of sessions (N) for each condition is reported in Table 1.

| Monkey | Task | Velocity component | R<sup>2</sup> (median [IQR]) |  | MSE (median [IQR]) |  | Combine p-value |  |
| --- | --- | --- | --- | --- | --- | --- | --- | --- |
|  |  |  | Linear | Non-linear | Linear | Non-linear | Linear | Non-linear |
| Monkey 1 | Center-out | v | 9.67 x 10<sup>-5</sup><br>[1.15 x 10<sup>-4</sup>] | 1.64 x 10<sup>-3</sup><br>[0.76 x 10<sup>-3</sup>] | 2.17 [0.2] | 2.16 [0.2] | 0.43 | 1.62 x 10<sup>-5</sup> |
|  |  | v<sub>x</sub> | 6.67 x 10<sup>-4</sup><br>[2.80 x 10<sup>-3</sup>] | 0.75 x 10<sup>-3</sup><br>[3.90 x 10<sup>-3</sup>] | 2.07 [0.49] | 2.06 [0.49] | 1.17 x 10<sup>-11</sup> | 2.53 x 10<sup>-5</sup> |
|  |  | v<sub>z</sub> | 4.19 x 10<sup>-5</sup><br>[7.61 x 10<sup>-4</sup>] | 1.84 x 10<sup>-3</sup><br>[9.34 x 10<sup>-3</sup>] | 1.64 [0.04] | 1.63 [0.04] | 4.85 x 10<sup>-3</sup> | 4.38 x 10<sup>-6</sup> |
|  | Continuous Navigation | v | 7.98 x 10<sup>-4</sup><br>[1.57 x 10<sup>-3</sup>] | 1.05 x 10<sup>-3</sup><br>[1.51 x 10<sup>-3</sup>] | 0.82 [0.23] | 0.82 [0.23] | 4.66 x 10<sup>-9</sup> | 1.48 x 10<sup>-2</sup> |
|  |  | v<sub>x</sub> | 6.0 x 10<sup>-3</sup><br>[22.9 x 10<sup>-3</sup>] | 10.9 x 10<sup>-3</sup><br>[25.6x 10<sup>-3</sup>] | 0.43 [0.06] | 0.43 [0.06] | 5.46 x 10<sup>-158</sup> | 2.44 x 10<sup>-42</sup> |
|  |  | v<sub>z</sub> | 1.15x 10<sup>-3</sup><br>[2.47 x 10<sup>-3</sup>] | 1.52 x 10<sup>-3</sup><br>[2.31 x 10<sup>-3</sup>] | 0.79 [0.2] | 0.79 [0.2] | 2.99 x 10<sup>-24</sup> | 0.09 |
| Monkey 2 | Center-out | v | 9.47 x 10<sup>-3</sup><br>[1.98 x 10<sup>-2</sup>] | 1.68 x 10<sup>-2</sup><br>[2.59 x 10<sup>-2</sup>] | 1.13 [0.18] | 112 [0.19] | 1.27 x 10<sup>-150</sup> | 2.07 x 10<sup>-95</sup> |
|  |  | v<sub>x</sub> | 6.62 x 10<sup>-3</sup><br>[2.81 x 10<sup>-2</sup>] | 1.69 x 10<sup>-2</sup><br>[3.97x 10<sup>-2</sup>] | 2.64 [0.71] | 2.59 [0.72] | 0.0 | 2.54 x 10<sup>-105</sup> |
|  |  | v<sub>z</sub> | 5.37x 10<sup>-3</sup><br>[9.44 x 10<sup>-3</sup>] | 8.30 x 10<sup>-3</sup><br>[1.37 x 10<sup>-2</sup>] | 0.90 [0.21] | 0.90 [0.21] | 6.96 x 10<sup>-72</sup> | 5.72 x 10<sup>-51</sup> |
|  | Continuous Navigation | v | 2.88 x 10<sup>-3</sup><br>[6.83 x 10<sup>-3</sup>] | 6.23 x 10<sup>-3</sup><br>[1.57 x 10<sup>-3</sup>] | 0.55 [0.59] | 0.54 [0.59] | 5.09 x 10<sup>-42</sup> | 5.38 x 10<sup>-13</sup> |
|  |  | v<sub>x</sub> | 8.32 x 10<sup>-3</sup><br>[1.06 x 10<sup>-2</sup>] | 1.02 x 10<sup>-2</sup><br>[1.34 x 10<sup>-2</sup>] | 1.23 [0.64] | 1.22 [0.64] | 1.77 x 10<sup>-129</sup> | 9.13 x 10<sup>-24</sup> |
|  |  | v<sub>z</sub> | 1.02 x 10<sup>-3</sup><br>[2.49 x 10<sup>-3</sup>] | 2.97 x 10<sup>-3</sup><br>[3.12 x 10<sup>-3</sup>] | 1.58 [0.65] | 1.58 [0.64] | 2.44 x 10<sup>-30</sup> | 4.44 x 10<sup>-11</sup> |
|  | 3D Center-out | v | 7.93 x 10<sup>-3</sup><br>[1.65 x 10<sup>-2</sup>] | 1.25 x 10<sup>-2</sup><br>[12.51 x 10<sup>-2</sup>] | 1.05 [0.39] | 1.05 [0.39] | 1.30 x 10<sup>-172</sup> | 4.98 x 10<sup>-98</sup> |
|  |  | v<sub>x</sub> | 3.56 x 10<sup>-2</sup><br>[5.42 x 10<sup>-2</sup>] | 4.42 x 10<sup>-2</sup><br>[4.34 x 10<sup>-2</sup>] | 2.78 [0.76] | 2.74 [0.72] | 0.0 | 4.52 x 10<sup>-184</sup> |
|  |  | v<sub>y</sub> | 1.09 x 10<sup>-2</sup><br>[2.45 x 10<sup>-2</sup>] | 1.45 x 10<sup>-2</sup><br>[2.27 x 10<sup>-2</sup>] | 0.06 [0.02] | 0.06 [0.02] | 6.81 x 10<sup>-220</sup> | 2.29 x 10<sup>-62</sup> |
|  |  | v<sub>z</sub> | 4.55 x 10<sup>-3</sup><br>[6.30 x 10<sup>-3</sup>] | 5.50 x 10<sup>-3</sup><br>[1.26 x 10<sup>-2</sup>] | 0.74 [0.36] | 0.73 [0.36] | 1.81 x 10<sup>-102</sup> | 9.70 x 10<sup>-43</sup> |
|  | Respawn | v | 6.59 x 10<sup>-3</sup><br>[1.95 x 10<sup>-2</sup>] | 8.44 x 10<sup>-3</sup><br>[1.64 x 10<sup>-2</sup>] | 0.83 [0.24] | 0.82 [0.24] | 7.74 x 10<sup>-181</sup> | 3.03 x 10<sup>-21</sup> |
|  |  | v<sub>x</sub> | 3.08 x 10<sup>-2</sup><br>[8.41 x 10<sup>-2</sup>] | 5.45 x 10<sup>-2</sup><br>[7.97x 10<sup>-2</sup>] | 2.26 [0.76] | 2.26 [0.76] | 0.0 | 1.48 x 10<sup>-115</sup> |
|  |  | v<sub>z</sub> | 1.70x 10<sup>-3</sup><br>[1.04 x 10<sup>-2</sup>] | 2.58x 10<sup>-3</sup><br>[1.72 x 10<sup>-2</sup>] | 0.70 [0.06] | 0.70 [0.067] | 1.02 x 10<sup>-111</sup> | 1.00 x 10<sup>-36</sup> |
|  | Obstacle | v | 6.12 x 10<sup>-3</sup><br>[9.57 x 10<sup>-3</sup>] | 9.91 x 10<sup>-3</sup><br>[8.34x 10<sup>-3</sup>] | 1.08 [0.26] | 1.08 [0.26] | 1.33 x 10<sup>-68</sup> | 6.57 x 10<sup>-19</sup> |
|  |  | v<sub>x</sub> | 1.60 x 10<sup>-2</sup><br>[2.22 x 10<sup>-2</sup>] | 3.13 x 10<sup>-2</sup><br>[4.32 x 10<sup>-2</sup>] | 2.58 [0.30] | 2.57 [0.32] | 2.07 x 10<sup>-206</sup> | 1.88 x 10<sup>-133</sup> |
|  |  | v<sub>z</sub> | 2.22 x 10<sup>-3</sup><br>[3.36 x 10<sup>-3</sup>] | 3.36 x 10<sup>-3</sup><br>[4.48 x 10<sup>-3</sup>] | 0.85 [0.10] | 0.85 [0.10] | 6.17 x 10<sup>-42</sup> | 5.02 x 10<sup>-7</sup> |

**Table S4.**
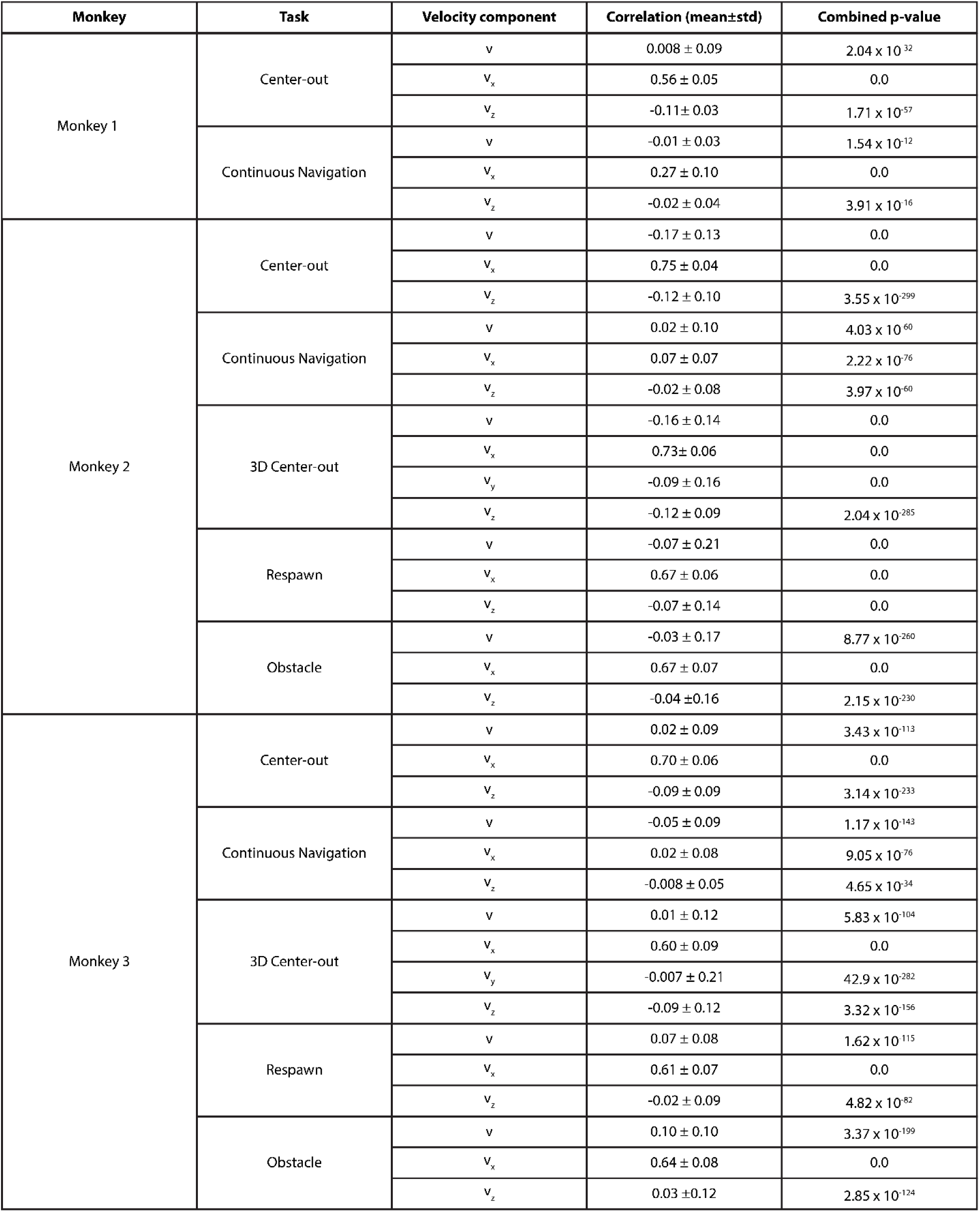
Spearman correlation and combined p-value between x-component of eye movement and different velocity components for Monkey 1, 2 and 3, for various tasks. The mean ± standard deviation Spearman correlation and combined p-value (Fisher’s method) across all sessions. N (number of sessions) is listed in Table 1.

| Monkey | Task | Velocity component | Correlation (mean±std) | Combined p-value |
| --- | --- | --- | --- | --- |
| Monkey 1 | Center-out | v | 0.008 ± 0.09 | 2.04 × 10<sup>-32</sup> |
|  |  | v<sub>x</sub> | 0.56 ± 0.05 | 0.0 |
|  |  | v<sub>z</sub> | -0.11 ± 0.03 | 1.71 × 10<sup>-57</sup> |
|  | Continuous Navigation | v | -0.01 ± 0.03 | 1.54 × 10<sup>-12</sup> |
|  |  | v<sub>x</sub> | 0.27 ± 0.10 | 0.0 |
|  |  | v<sub>z</sub> | -0.02 ± 0.04 | 3.91 × 10<sup>-16</sup> |
| Monkey 2 | Center-out | v | -0.17 ± 0.13 | 0.0 |
|  |  | v<sub>x</sub> | 0.75 ± 0.04 | 0.0 |
|  |  | v<sub>z</sub> | -0.12 ± 0.10 | 3.55 × 10<sup>-299</sup> |
|  | Continuous Navigation | v | 0.02 ± 0.10 | 4.03 × 10<sup>-60</sup> |
|  |  | v<sub>x</sub> | 0.07 ± 0.07 | 2.22 × 10<sup>-76</sup> |
|  |  | v<sub>z</sub> | -0.02 ± 0.08 | 3.97 × 10<sup>-60</sup> |
|  | 3D Center-out | v | -0.16 ± 0.14 | 0.0 |
|  |  | v<sub>x</sub> | 0.73 ± 0.06 | 0.0 |
|  |  | v<sub>y</sub> | -0.09 ± 0.16 | 0.0 |
|  |  | v<sub>z</sub> | -0.12 ± 0.09 | 2.04 × 10<sup>-285</sup> |
|  | Respawn | v | -0.07 ± 0.21 | 0.0 |
|  |  | v<sub>x</sub> | 0.67 ± 0.06 | 0.0 |
|  |  | v<sub>z</sub> | -0.07 ± 0.14 | 0.0 |
|  | Obstacle | v | -0.03 ± 0.17 | 8.77 × 10<sup>-260</sup> |
|  |  | v<sub>x</sub> | 0.67 ± 0.07 | 0.0 |
|  |  | v<sub>z</sub> | -0.04 ± 0.16 | 2.15 × 10<sup>-230</sup> |
| Monkey 3 | Center-out | v | 0.02 ± 0.09 | 3.43 × 10<sup>-113</sup> |
|  |  | v<sub>x</sub> | 0.70 ± 0.06 | 0.0 |
|  |  | v<sub>z</sub> | -0.09 ± 0.09 | 3.14 × 10<sup>-233</sup> |
|  | Continuous Navigation | v | -0.05 ± 0.09 | 1.17 × 10<sup>-143</sup> |
|  |  | v<sub>x</sub> | 0.02 ± 0.08 | 9.05 × 10<sup>-76</sup> |
|  |  | v<sub>z</sub> | -0.008 ± 0.05 | 4.65 × 10<sup>-34</sup> |
|  | 3D Center-out | v | 0.01 ± 0.12 | 5.83 × 10<sup>-104</sup> |
|  |  | v<sub>x</sub> | 0.60 ± 0.09 | 0.0 |
|  |  | v<sub>y</sub> | -0.007 ± 0.21 | 42.9 × 10<sup>-282</sup> |
|  |  | v<sub>z</sub> | -0.09 ± 0.12 | 3.32 × 10<sup>-156</sup> |
|  | Respawn | v | 0.07 ± 0.08 | 1.62 × 10<sup>-115</sup> |
|  |  | v<sub>x</sub> | 0.61 ± 0.07 | 0.0 |
|  |  | v<sub>z</sub> | -0.02 ± 0.09 | 4.82 × 10<sup>-82</sup> |
|  | Obstacle | v | 0.10 ± 0.10 | 3.37 × 10<sup>-199</sup> |
|  |  | v<sub>x</sub> | 0.64 ± 0.08 | 0.0 |
|  |  | v<sub>z</sub> | 0.03 ± 0.12 | 2.85 × 10<sup>-124</sup> |

**Table S5.**
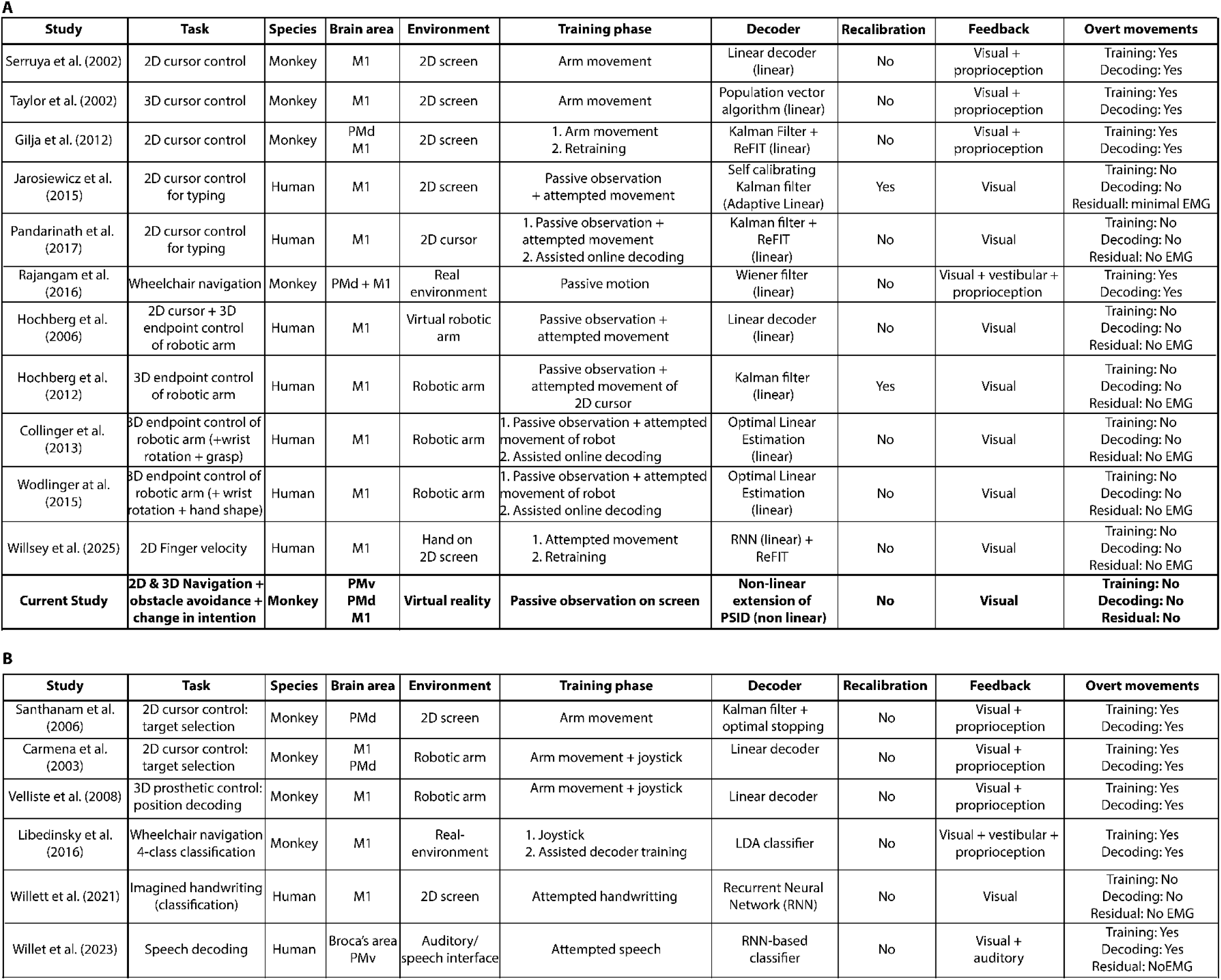
Intracortical motor BCI studies using velocity, position, classification, or language decoding. A. Summary of representative intracortical BCI studies that employ continuous velocity decoding in motor tasks with online feedback. For each study, we report task type, species, environment, training paradigm, decoder architecture, recalibration, feedback modality, and the presence of overt movement or EMG monitoring. Our current study (highlighted) is the only one to feature nonlinear decoding of 3D navigation using passive observation alone, without decoder recalibration or overt movement. Success rates and performance metrics vary across studies and are not directly comparable due to differences in task design, autonomy levels, and success criteria. Key performance indicators are provided below for contextual understanding: Serruya (2002): not reported; Taylor (2004): 49%; Gilja (2012): 100% (target window enlarged for method comparison); Hochberg (2006): 73–95%; Hochberg (2012): 69–96%; Collinger (2013): 85% (7 DOF); Wodlinger (2015): 70% (10 DOF); Jarosiewicz (2015): typing – 12 characters/min; Pandarinath (2016): typing – 12–39 characters/min. **B** Complementary studies using position decoding, classification, or language models rather than continuous velocity decoding. While not directly comparable, these studies illustrate the diversity of intracortical decoding strategies across motor, cognitive, and speech-related tasks.

